# Defining the transcriptional and post-transcriptional landscapes of *Mycobacterium smegmatis* in aerobic growth and oxygen limitation

**DOI:** 10.1101/423392

**Authors:** M. Carla Martini, Ying Zhou, Huaming Sun, Scarlet S. Shell

## Abstract

The ability of *Mycobacterium tuberculosis* to infect, proliferate, and survive during long periods in the human lungs largely depends on the rigorous control of gene expression. Transcriptome-wide analyses are key to understanding gene regulation on a global scale. Here, we combine 5’-end-directed libraries with RNAseq expression libraries to gain insight into the transcriptome organization and post-transcriptional mRNA cleavage landscape in mycobacteria during log phase growth and under hypoxia, a physiologically relevant stress condition. Using the model organism *Mycobacterium smegmatis*, we identified 6,090 transcription start sites (TSSs) with high confidence during log phase growth, of which 67% were categorized as primary TSSs for annotated genes, and the remaining were classified as internal, antisense or orphan, according to their genomic context. Interestingly, over 25% of the RNA transcripts lack a leader sequence, and of the coding sequences that do have leaders, 53% lack a strong consensus Shine-Dalgarno site. This indicates that like *M. tuberculosis*, *M. smegmatis* can initiate translation through multiple mechanisms. Our approach also allowed us to identify over 3,000 RNA cleavage sites, which occur at a novel sequence motif. The cleavage sites show a positional bias toward mRNA regulatory regions, highlighting the importance of post-transcriptional regulation in gene expression. We show that in low oxygen, a condition associated with the host environment during infection, mycobacteria change their transcriptomic profiles and endonucleolytic RNA cleavage is markedly reduced, suggesting a mechanistic explanation for previous reports of increased mRNA half-lives in response to stress. In addition, a number of TSSs were triggered in hypoxia, 56 of which contain the binding motif for the sigma factor SigF in their promoter regions. This suggests that SigF makes direct contributions to transcriptomic remodeling in hypoxia-challenged mycobacteria. Our results show that *M. smegmatis* and *M. tuberculosis* share a large number of similarities at the transcriptomic level, suggesting that similar regulatory mechanisms govern both species.

## Introduction

Tuberculosis is a disease of global concern caused by *Mycobacterium tuberculosis* (Mtb). This pathogen has the ability to infect the human lungs and survive there for long periods, often by entering into non-growing states. During infection, Mtb must overcome a variety of stressful conditions, including nutrient starvation, low pH, oxygen deprivation and the presence of reactive oxygen species. Consequently, the association of Mtb with its host and the adaptation to the surrounding environment requires rigorous control of gene expression.

As the slow growth rate and pathogenicity of Mtb present logistical challenges in the laboratory, many aspects of its biology have been studied in other mycobacterial species. One of the most widely used models is mycobacteria is *Mycobacterium smegmatis*, a non-pathogenic fast-growing bacterium that shares substantial genomic similarity with Mtb. A PubMed search for “*Mycobacterium smegmatis*” returns 3,907 publications, reflecting the sizable body of published work involving this model organism. While there are marked differences between the genomes, such as the highly represented PE/PPE-like gene category and other virulence factors present in Mtb and poorly represented or absent in *M. smegmatis*, these organisms have at least 2,117 orthologous genes (Prasanna & Mehra, 2013) making *M. smegmatis* a viable model to address questions about the fundamental biology of mycobacteria. Indeed, studies using *M. smegmatis* have revealed key insights into relevant aspects of Mtb biology including the Sec and ESX secretion systems involved in transport of virulence factors (Coros *et al.*, 2008, Rigel *et al.*, 2009), bacterial survival during anaerobic dormancy (Dick *et al.*, 1998, Bagchi *et al.*, 2002, Trauner *et al.*, 2012, Pecsi *et al.*, 2014) and the changes induced during nutrient starvation (Elharar *et al.*, 2014, Wu *et al.*, 2016, Hayashi *et al.*, 2018). However, the similarities and differences between *M. smegmatis* and *M. tuberculosis* at the level of transcriptomic organization have not been comprehensively reported.

Identification of transcription start sites (TSSs) is an essential step towards understanding how bacteria organize their transcriptomes and respond to changing environments. Genome-wide TSS mapping studies have been used to elucidate the general transcriptomic features in many bacterial species, leading to the identification of promoters, characterization of 5’ untranslated regions (5’ UTRs), identification of RNA regulatory elements and transcriptional changes in different environmental conditions (examples include (Albrecht *et al.*, 2009, Mitschke *et al.*, 2011, Cortes *et al.*, 2013, Schlüter *et al.*, 2013, Dinan *et al.*, 2014, Ramachandran *et al.*, 2014, Sass *et al.*, 2015, Shell *et al.*, 2015, Thomason *et al.*, 2015, Berger *et al.*, 2016, Čuklina *et al.*, 2016, D’arrigo *et al.*, 2016, Heidrich *et al.*, 2017, Li *et al.*, 2017). To date, two main studies have reported the transcriptomic landscape in Mtb during exponential growth and carbon starvation (Cortes *et al.*, 2013, Shell *et al.*, 2015). These complementary studies revealed that, unlike most bacteria, a substantial percentage (~25%) of the transcripts are leaderless, lacking a 5’ UTR and consequently a Shine-Dalgarno ribosome-binding site. In addition, a number of previously unannotated ORFs encoding putative small proteins were found (Shell *et al.*, 2015), showing that the transcriptional landscape can be more complex than predicted by automated genome annotation pipelines. Thus, TSS mapping is a powerful tool to gain insight into transcriptomic organization and identify novel genes. Less is known about the characteristics of the *M. smegmatis* transcriptome. A recent study reported a number of *M. smegmatis* TSSs in normal growth conditions (Li *et al.* 2017). However, this work was limited to identification of primary gene-associated TSSs and lacked of an analysis of internal and antisense TSSs, as well as characterization of promoter regions and other relevant transcriptomic features. In addition, Potgieter and collaborators (Potgieter *et al.*, 2016) validated a large number of annotated ORFs using proteomics and were able to identify 63 previously unannotated leaderless ORFs.

To achieve a deeper characterization of the *M. smegmatis* transcriptional landscape, we combined 5’-end-mapping and RNAseq expression profiling under two different growth conditions. Here we present an exhaustive analysis of *M. smegmatis* transcriptome during exponential growth and hypoxia. Unlike most transcriptome-wide TSS analyses, our approach allowed us to study not only the transcriptome organization in different conditions, but also the frequency and distribution of RNA cleavage sites on a genome wide scale. Whereas regulation at the transcriptional level is assumed to be the main mechanism that modulates gene expression in bacteria, post-transcriptional regulation is a key step in the control of gene expression and has been implicated in the response to host conditions and virulence in various bacterial pathogens (Kulesekara *et al.*, 2006, Mraheil *et al.*, 2011, Heroven *et al.*, 2012, Jurėnaitė *et al.*, 2013, Schifano *et al.*, 2013, Holmqvist *et al.*, 2016). Some regulatory mechanisms including small non-coding RNAs, RNases, Toxin-Antitoxin (TA) systems, RNA-binding proteins, and riboswitches have been described in mycobacteria ((Fields & Switzer, 2007, Warner *et al.*, 2007, Sala *et al.*, 2008, DiChiara *et al.*, 2010, McKenzie *et al.*, 2012, Winther *et al.*, 2016) and others), emphasizing the importance of post-transcriptional regulation. Here we show that RNA cleavage decreases during adaptation to hypoxia, suggesting that RNA cleavage may be a refinement mechanism contributing to the regulation of gene expression in harsh conditions.

## Materials and Methods

### Strains and growth conditions used in this study

*M. smegmatis* strain mc^2^155 was grown in Middlebrook 7H9 supplemented with ADC (Albumin Dextrose Catalase, final concentrations 5 g/L bovine serum albumin fraction V, 2 g/L dextrose, 0.85 g/L sodium chloride, and 3 mg/L catalase), 0.2% glycerol and 0.05% Tween 80. For the exponential phase experiment (Dataset 1), 50 ml conical tubes containing 5 ml of 7H9 were inoculated with *M. smegmatis* to have an initial OD=0.01. Cultures were grown at 37°C and 250 rpm. Once cultures reached an OD of 0.7 – 0.8, they were frozen in liquid nitrogen and stored at -80°C until RNA purification. For hypoxia experiments (Dataset 2), the Wayne model (Wayne & Hayes, 1996) was implemented. Briefly, 60 ml serum bottles (Wheaton) were inoculated with 36.5 ml of *M. smegmatis* culture with an initial OD=0.01. The bottles were sealed with rubber caps (Wheaton, W224100-181 Stopper, 20mm) and aluminum caps (Wheaton, 20 mm aluminum seal) and cultures were grown at 37 °C and 125 rpm to generate hypoxic conditions. Samples were taken at an early hypoxia stage (15 hours) and at a late hypoxia stage (24 hours). These time points were experimentally determined according to growth curves experiments (see **Figure S1**). 15 ml of each culture were sampled and frozen immediately in liquid nitrogen until RNA extraction.

### RNA extraction

RNA was extracted as follows: frozen cultures stored at -80°C were thawed on ice and centrifuged at 4,000 rpm for 5 min at 4 °C. The pellets were resuspended in 1 ml Trizol (Life Technologies) and placed in tubes containing Lysing Matrix B (MP Bio). Cells were lysed by bead-beating twice for 40 sec at 9 m/sec in a FastPrep 5G instrument (MP Bio). 300 μl chloroform was added and samples were centrifuged for 15 minutes at 4,000 rpm at 4°C. The aqueous phase was collected and RNA was purified using Direct-Zol RNA miniprep kit (Zymo) according to the manufacturer’s instructions. Samples were then treated with DNase Turbo (Ambion) for one hour and purified with a RNA Clean & Concentrator-25 kit (Zymo) according to the manufacturer’s instructions. RNA integrity was checked on 1% agarose gels and concentrations were determined using a Nanodrop instrument. Prior to library construction, 5 μg RNA was used for rRNA depletion using Ribo-Zero rRNA Removal Kit (Illumina) according to the manufacturer’s instructions.

### Construction of 5’-end-mapping libraries

After rRNA depletion, RNA samples from each biological replicate were split in three, in order to generate two 5’-end differentially treated libraries and one RNAseq expression library (next section). RNA for library 1 (“converted” library) was treated either with RNA 5’ pyrophosphohydrolase RPPH (NEB) (exponential phase experiment, Dataset 1), or with 5’ polyphosphatase (Epicentre) (hypoxia experiment, Dataset 2), in order to remove the native 5’ triphosphates of primary transcripts, whereas RNA for Library 2 (“non-converted” library) was subject to mock treatment. Thus, the converted libraries capture both 5’ triphosphates (converted to monophosphates) and native 5’ monophosphate transcripts, while non-converted libraries capture only native 5’ monophosphates (see scheme in **Figure S2.A**). Library construction was performed as described by Shell et al (Shell *et al.*, 2015). A detailed scheme showing the workflow of 5’-end libraries construction, the primers and adapters used in each step, and modifications to the protocol are shown in **Figure S2.B**.

### Construction of RNAseq expression libraries

One third of each rRNA-depleted RNA sample was used to construct RNAseq expression libraries. KAPA stranded RNA-Seq library preparation kit and NEBNext Ultra RNA library prep kit for Illumina (NEB) were used for Dataset 1 and Dataset 2, respectively, according to manufacturer’s instructions. The following major modifications were introduced into the protocols: *i)* For RNA fragmentation, in order to obtain fragments around 300 nt, RNA was mixed with the corresponding buffer and placed at 85°C for 6 minutes (Dataset 1), or at 94°C for 12 minutes (Dataset 2). *ii)* For library amplification, 10 or 19-23 PCR cycles were used for Dataset 1 and Dataset 2, respectively. The number of cycles was chosen according to the amount of cDNA obtained for each sample. After purification, DNA concentration was measured in a Qubit instrument before sequencing.

### Libraries sequencing and quality assessment

For 5’-end-mapping libraries from Dataset 1, Illumina MiSeq paired-end sequencing producing 100 nt reads was used. For 5’ end directed libraries from Dataset 2 as well as for all expression libraries, Illumina HiSeq 2000 paired-end sequencing producing 50 nt reads was used. Sequencing was performed at the UMass Medical School Deep Sequencing Core Facility. Quality of the generated fastq files was checked using FastQC.

### Identification of 5’ ends and discrimination between transcription start sites (TSSs) and cleavage sites (CSs)

Paired-end reads generated from 5’-end-directed libraries were mapped to *M. smegmatis* mc^2^155 NC_008596 reference genome. In order to reduce noise from the imprecision of transcriptional initiation, only the coordinate with the highest coverage in each 5 nt window was used for downstream analyses. For read filtering, different criteria were used for the 2 datasets according to the library depth and quality (see **Figure S3**). In order to discriminate between TSSs and CSs, the ratio of the coverage in converted/non-converted libraries for each detected 5’ end was calculated. To focus our analyses on the 5’ends that are relatively abundant in their local genomic context, we employed a filter based on the ratio of 5’ end coverage to expression library coverage in the preceding 100nt. 5’ ends for which this ratio was ≤0.05 were removed. After this filter, 15,720 5’ ends remained and were further analyzed using a Gaussian mixture modelling to differentiate TSSs from CSs with a high confidence in Dataset 1 (**Figure 1A**). For this analysis, we used the iterative expectation maximization (EM) algorithm in the mixtools package (Benaglia *et al.*, 2009) for R (version 1.1.0) to fit the mixture distributions.

**Figure 1.**
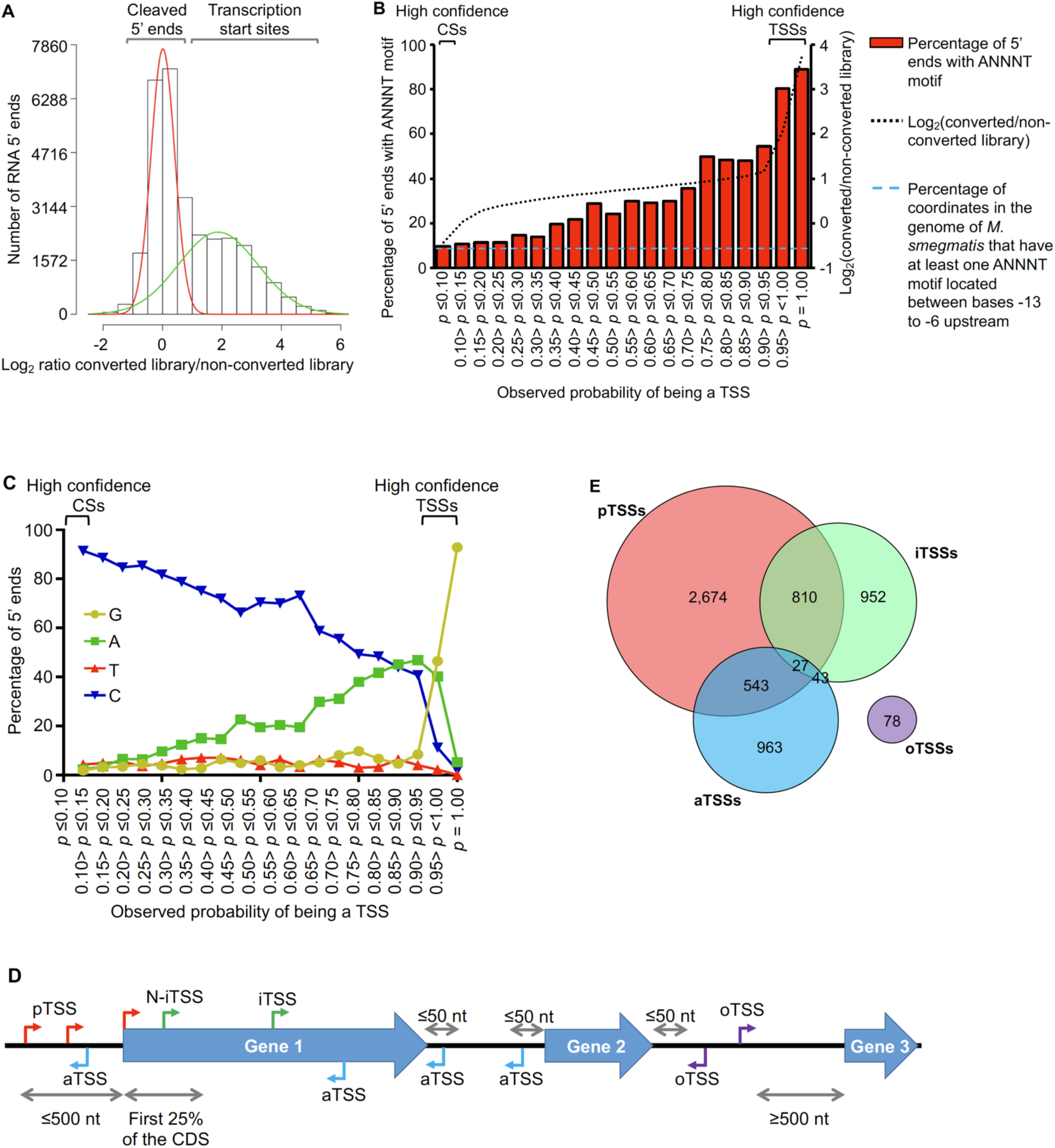
Mapping and categorization of transcription start sites in *M. smegmatis*. **A)** Diagram showing the ratios of coverage in the converted/non-converted libraries for each coordinate. Gaussian mixture modeling was used to discriminate between TSSs and CSs. For this analysis, the 15,720 coordinates from Dataset 1 were used. **B)** Abundance of the ANNNT promoter motif located between bases -13 to -6 upstream of the 15,720 coordinates. The light blue dashed line indicates the percentage of coordinates in the genome of *M. smegmatis* that have at least one ANNNT motif located between bases -13 to -6 upstream (9.7%). **C)** Base frequency at the +1 position among the 15,720 5’ ends from Dataset 1. Probabilities of a 5’ end being a TSS were calculated by Gaussian Mixture modeling. **D)** Categories for TSS annotation based on the genomic context. TSSs were classified according to their relative position to genes as primary (red), internal (green), antisense (light blue) and orphan (violet). **E)** Distribution of TSSs among the different categories.

### Analysis of expression libraries

Reads were aligned to NC_008596 reference genome using Burrows-Wheeler Aligner (Li & Durbin, 2009). For comparison of gene expression levels according to presence or absence of Shine-Dalgarno sequences, RPKMs were calculated for all genes. The DEseq2 pipeline was used to evaluate the changes in gene expression in hypoxia (Love *et al.*, 2014).

### Transcription start sites categorization

For analysis in **Figure 1D**, TSSs were classified as follows: those coordinates located ≤ 500 bp upstream from an annotated gene were considered to be primary TSSs (pTSS). Coordinates located within an annotated gene were classified as internal (iTSS) or N-associated internal TSSs (N-iTSSs) if they were located within the first 25% of the annotated coding sequence. N-iTSSs were considered for reannotation as a pTSSs only if their associated gene lacked a pTSS. TSSs located on the antisense strand of a coding sequence, 5’ UTR, or 3’ UTR were considered as antisense (aTSS). 5’ UTRs boundaries were assigned after assignment of pTSSs to genes annotated in the reference genome NC_008596. When a gene had more than one pTSS, the longest of the possible 5’ UTRs was used for assignment of aTSSs. In the case of genes for which we did not identify a pTSS, we considered a hypothetical leader sequence of 50 bp for assignment of aTSSs. For assignment of aTSSs in 3’ UTRs, we arbitrarily considered a sequence of 50 bp downstream the stop codon of the gene to be the 3’ UTR. Finally, TSSs not belonging to any of the above-mentioned categories were classified as orphan (oTSSs).

### Operon prediction

Adjacent genes with the same orientation were considered to be co-transcribed if there were at least 5 spanning reads between the upstream and the downstream gene in at least one of the replicates in the expression libraries from Dataset 1. After this filtering, a downstream gene was excluded from the operon if: 1) it had a TSS ≤ 500 bp upstream the annotated start codon on the same strand, and/or 2) had a TSS within the first 25% of the gene on the same strand, and/or 3) the upstream gene had a TSS within the last 50-100% of the coding sequence. Finally, the operon was assigned only if the first gene had a primary TSS with a confidence ≥ 95% according to the Gaussian mixture modeling.

### Cleavage sites categorization

For CS categorization in **Figure 4D**, we stablished stringent criteria in order to determine the frequency of CSs in each location category relative to the amount of the genome comprising that location category. For 3’ UTR regions, we considered only CSs that were located between 2 convergent genes. To assess frequency relative to the whole genome, we considered the sum of all regions located between two convergent genes. For 5’ UTRs we considered all CSs located between 2 divergent genes, and the sum of all leader lengths for genes having a pTSS whose upstream gene is in the opposite strand (divergent) determined in this study was used for assessing relative frequency. For 5’ ends corresponding to cleavages between co-transcribed genes we used the operon structures determined in this study, and the sum of all their intergenic regions was used for assessing relative frequency. Finally, for CSs located within coding sequences all genes were considered, as all of them produced reads in the expression libraries. The sum of all coding sequences in NC_008596 genome was used for assessing relative frequency, after subtracting overlapping regions to avoid redundancy.

## Results

### 1. Mapping, annotation and categorization of transcription start sites

In order to study the transcriptome structure of *M. smegmatis*, RNAs from triplicate cultures in exponential phase were used to construct 5’ end mapping libraries (Dataset 1, D1) according to our previously published methodologies (Shell *et al.*, 2015, Shell *et al.*, 2015) with minor modifications. Briefly, our approach relies on comparison of adapter ligation frequency in a dephosphorylated (converted) library and an untreated (non-converted) library for each sample. The converted libraries capture both 5’ triphosphate and native 5’ monophosphate-bearing transcripts, while the non-converted libraries capture only native 5’ monophosphate-bearing transcripts (**Figure S2**). Thus, assessing the ratios of read counts in the converted/non-converted libraries permits discrimination between 5’ triphosphate ends (primary transcripts from transcription start sites) and 5’ monophosphate ends (cleavage sites). By employing a Gaussian mixture modeling analysis (**Figure 1A**) we were able to identify 5,552 TSSs in *M. smegmatis* with an observed probability of being a TSS ≥0.95 (high confidence TSSs, **Table S1**). A second filtering method allowed us to obtain 222 additional TSSs from Dataset 1 (**Figure S3**). A total of 5,774 TSSs were therefore obtained from Dataset 1. In addition, data from separate libraries constructed as controls for the hypoxia experiment (Dataset 2, D2) in Section 8 were also included in this analysis to obtain TSSs. After noise filtering (**Figure S3**), 4,736 TSSs from D2 were identified. The union of the two datasets yielded a total of 6,090 non-redundant high confidence TSSs, of which 4,420 were detected in both datasets (**Figure S4**, **Table S1**).

Although not all 5’ ends could be classified with the Gaussian mixture modeling, we were able to assign 57% of the 5’ ends in Dataset 1 to one of the two 5’ end populations with high confidence (5,552 TSSs and 3,344 CSs). To validate the reliability of the Gaussian mixture modeling used to classify 5’ ends, we performed two additional analyses. First, according to previous findings in Mtb (Cortes *et al.*, 2013) and other well studied bacteria (Sass *et al.*, 2015, Berger *et al.*, 2016, Čuklina *et al.*, 2016, D’arrigo *et al.*, 2016), we anticipated that TSSs should be enriched for the presence of the ANNNT -10 promoter consensus motif in the region upstream. Evaluation of the presence of appropriately-spaced ANNNT sequences revealed that 5’ ends with higher probabilities of being TSSs are enriched for this motif, whereas for those 5’ ends with low probabilities of being TSSs (and thus high probabilities of being CSs) have ANNNT frequencies similar to that of the *M. smegmatis* genome as a whole (**Figure 1B**). Secondly, we predicted that TSSs should show enrichment for A and G nts at the +1 position, given the reported preference for bacterial RNA polymerases to initiate transcription with these nts (Lewis & Adhya, 2004, Mendoza-Vargas *et al.*, 2009, Mitschke *et al.*, 2011, Cortes *et al.*, 2013, Shell *et al.*, 2015, Thomason *et al.*, 2015, Berger *et al.*, 2016). Thus, we analyzed the base enrichment in the +1 position for the 5’ ends according the *p*-value in the Gaussian mixture modeling (**Figure 1C**). These results show a clear increase in the percentage of G and A bases in the position +1 as the probability of being a TSS increases, while the percentage of sequences having a C at +1 increases as the probability of being a TSS decreases. These two analyses show marked differences in the sequence contexts of TSSs and CSs and further validate the method used for categorization of 5’ ends.

To study the genome architecture of *M. smegmatis*, the 6,090 TSSs were categorized according to their genomic context (**Figure 1D and 1E**, **Table S1**). TSSs located ≤500 nt upstream of an annotated gene start codon in the *M. smegmatis* NC_008596 reference genome were classified as primary TSSs (pTSS). TSSs within annotated genes on the sense strand were denoted as internal (iTSS). When an iTSS was located in the first quarter of an annotated gene, it was sub-classified as N-terminal associated TSS (N-iTSS), and was further examined to determine if it should be considered a primary TSS (see below). TSSs located on the antisense strand either within a gene or within a 5’ UTR or 3’ UTR were grouped as antisense TSSs (aTSSs). Finally, TSSs located in non-coding regions that did not meet the criteria for any of the above categories were classified as orphan (oTSSs). When a pTSS also met the criteria for classification in another category, it was considered to be pTSS for the purposes of downstream analyses. A total of 4,054 distinct TSSs met the criteria to be classified as pTSSs for genes transcribed in exponential phase. These pTSSs were assigned to 3,043 downstream genes, representing 44% of the total annotated genes (**Table S2**). This number is lower than the total number of genes expressed in exponential phase, in large part due to the existence of polycistronic transcripts (see operon prediction below). Interestingly, 706 (23%) of these genes have at least two pTSSs and 209 (7%) have three or more, indicating that transcription initiation from multiple promoters is common in *M. smegmatis*.

A total of 995 iTSSs (excluding the iTSSs that were also classified as a pTSS of a downstream gene, see **Figure S5** for classification workflow) were identified in 804 (12%) of the annotated genes, indicating that transcription initiation within coding sequences is common in *M. smegmatis*. iTSSs are often considered to be pTSSs of downstream genes, to be spurious events yielding truncated transcripts, or to be consequences of incorrect gene start annotations. However, there is evidence supporting the hypothesis that iTSSs are functional and highly conserved among closely related bacteria (Shao *et al.*, 2014), highlighting their potential importance in gene expression.

We were also able to detect antisense transcription in 12.5 % of the *M. smegmatis* genes. Antisense transcription plays a role in modulation of gene expression by controlling transcription, RNA stability, and translation (Morita *et al.*, 2005, Kawano *et al.*, 2007, Andre *et al.*, 2008, Fozo *et al.*, 2008, Giangrossi *et al.*, 2010) and has been found to occur at different rates across bacterial genera, ranging from 1.3% of genes in *Staphylococcus aureus* to up to 46% of genes in *Helicobacter pylori* (Beaume *et al.*, 2010, Sharma *et al.*, 2010). Of the 1,006 aTSSs identified here (excluding those that were primarily classified as pTSSs), 881 are within coding sequences, 120 are within 5’ UTRs and 72 are located within 3’ UTRs (note that some aTSS are simultaneously classified in more than one of these three subcategories, **Figure S6**). While we expect that many of the detected antisense transcripts have biological functions, it is difficult to differentiate antisense RNAs with regulatory functions from transcriptional noise. In this regard, Lloréns-Rico and collaborators (Llorens-Rico & Cano, 2016) reported that most of the antisense transcripts detected using transcriptomic approaches are a consequence of transcriptional noise, arising at spurious promoters throughout the genome. To investigate the potential significance of the *M. smegmatis* aTSSs, we assessed the relative impact of each aTSS on local antisense expression levels by comparing the read depth upstream and downstream of each aTSS in our RNAseq expression libraries. We found 318 aTSSs for which expression coverage was ≥10-fold higher in the 100 nt window downstream of the TSS compared to the 100 nt window upstream (**Table S3**). Based on the magnitude of the expression occurring at these aTSS, we postulate that they could represent the 5’ ends of candidate functional antisense transcripts rather than simply products of spurious transcription. However, further work is needed to test this hypothesis. Finally, 78 oTSSs were detected across the *M. smegmatis* genome. These TSSs may be the 5’ ends of non-coding RNAs or mRNAs encoding previously unannotated ORFs.

Out of the 995 iTSSs identified, 457 were located within the first quarter of an annotated gene (N-iTSSs). In cases where there was no pTSS, we considered the possibility that the start codon of the gene was misannotated and the N-iTSS was in fact the primary TSS. Although we do not discount the possibility that functional proteins can be produced when internal transcription initiation occurs far downstream of the annotated start codon, we only considered N-iTSSs candidates for gene start reannotation when there was a start codon (ATG, GTG or TTG) in-frame with the annotated gene in the first 30% of the annotated sequence. In this way, we suggest re-annotations of the start codons of 213 coding sequences (see **Table S4**). These N-iTSSs were considered to be pTSSs (N-iTSSs → pTSSs) for all further analyses described in this work.

### 2. Operon prediction

To predict operon structure, we combined 5’ end libraries and RNAseq expression data. We considered two or more genes to be co-transcribed if (1) they had spanning reads that overlapped both the upstream and downstream gene in the expression libraries, (2) at least one TSS was detected in the 5’ end-directed libraries for the first gene of the operon, and (3) the downstream gene(s) lacked pTSSs and iTSSs (for more detail, see Materials and Methods). Thus, we were able to identify and annotate 294 operons with high confidence across the *M. smegmatis* genome (**Table S5**). These operons are between 2 and 4 genes in length and comprise a total of 638 genes. Our operon prediction methodology has some limitations. For example, operons not expressed in exponential growth phase could not be detected in our study. Furthermore, internal promoters within operons can exist, leading to either monocistronic transcripts or suboperons (Guell *et al.*, 2009, Paletta & Ohman, 2012, Skliarova *et al.*, 2012). We limited our operon predictions to genes that appear to be exclusively co-transcribed, excluding those cases in which an internal gene in an operon can be alternatively transcribed from an assigned pTSS. Finally, our analysis did not capture operons in which the first gene lacked a high-confidence pTSS. Despite these limitations, our approach allowed us to successfully identify new operons as well as previously described operons. Previously reported operons that were captured by our predictions included the *furA*-*katG* (MSMEG_6383-MSMEG_6384) operon involved in oxidative stress response (Milano *et al.*, 2001), the *vapB*-*vapC* (MSMEG_1283-MSMEG_1284) Toxin–Antitoxin module (Robson *et al.*, 2009) operon, and the *ClpP1*-*ClpP2* (MSMEG_4672-MSMEG_4673) operon involved in protein degradation (Raju *et al.*, 2012).

### 3. Characterization of *M. smegmatis* promoters reveals features conserved in *M. tuberculosis*

Most bacterial promoters have two highly conserved regions, the -10 and the -35, that interact with RNA polymerase via sigma factors. However, it was reported that the -10 region is necessary and sufficient for transcription initiation by the housekeeping sigma factor SigA in mycobacteria, and no SigA -35 consensus motifs were identified in previous studies (Cortes *et al.*, 2013, Newton-Foot & Gey van Pittius, 2013, Zhu *et al.*, 2017). to characterize the core promoter motifs in *M. smegmatis* on a global scale we analyzed the 50 bp upstream of the TSSs. We found that 4,833 of 6,090 promoters analyzed (79%) have an ANNNT motif located between positions -6 to -13 upstream the TSSs (**Figure 2A**). In addition, 63% of the promoters with ANNNT motifs have a thymidine preceding this sequence (**T**ANNNT). This motif is similar to that previously described in a transcriptome-wide analysis for Mtb (Cortes *et al.*, 2013) and for most bacterial promoters that are recognized by the σ^70^ housekeeping sigma factor (Ramachandran *et al.*, 2014, Sass *et al.*, 2015, Berger *et al.*, 2016, Čuklina *et al.*, 2016, D’arrigo *et al.*, 2016). However, no apparent bias towards specific bases in the NNN region was detected in our study or in Mtb, while in other bacteria such as *E. coli*, *S. enterica*, *B. cenocepacia*, *P. putida*, and *B. subtillis* an A/T preference was observed in this region (Jarmer *et al.*, 2001, Ramachandran *et al.*, 2014, Sass *et al.*, 2015, Berger *et al.*, 2016, D’arrigo *et al.*, 2016). We were unable to detect a consensus motif in the -35 region either using MEME server (Bailey *et al.*, 2015) or manually assessing the possible base-enrichment in the -35 region. Analysis of the sequences in the immediate vicinity of TSSs revealed that G and A are the most frequent bases at the +1 position, and C is considerably more abundant at -1 (**Figure 2B**).

**Figure 2.**
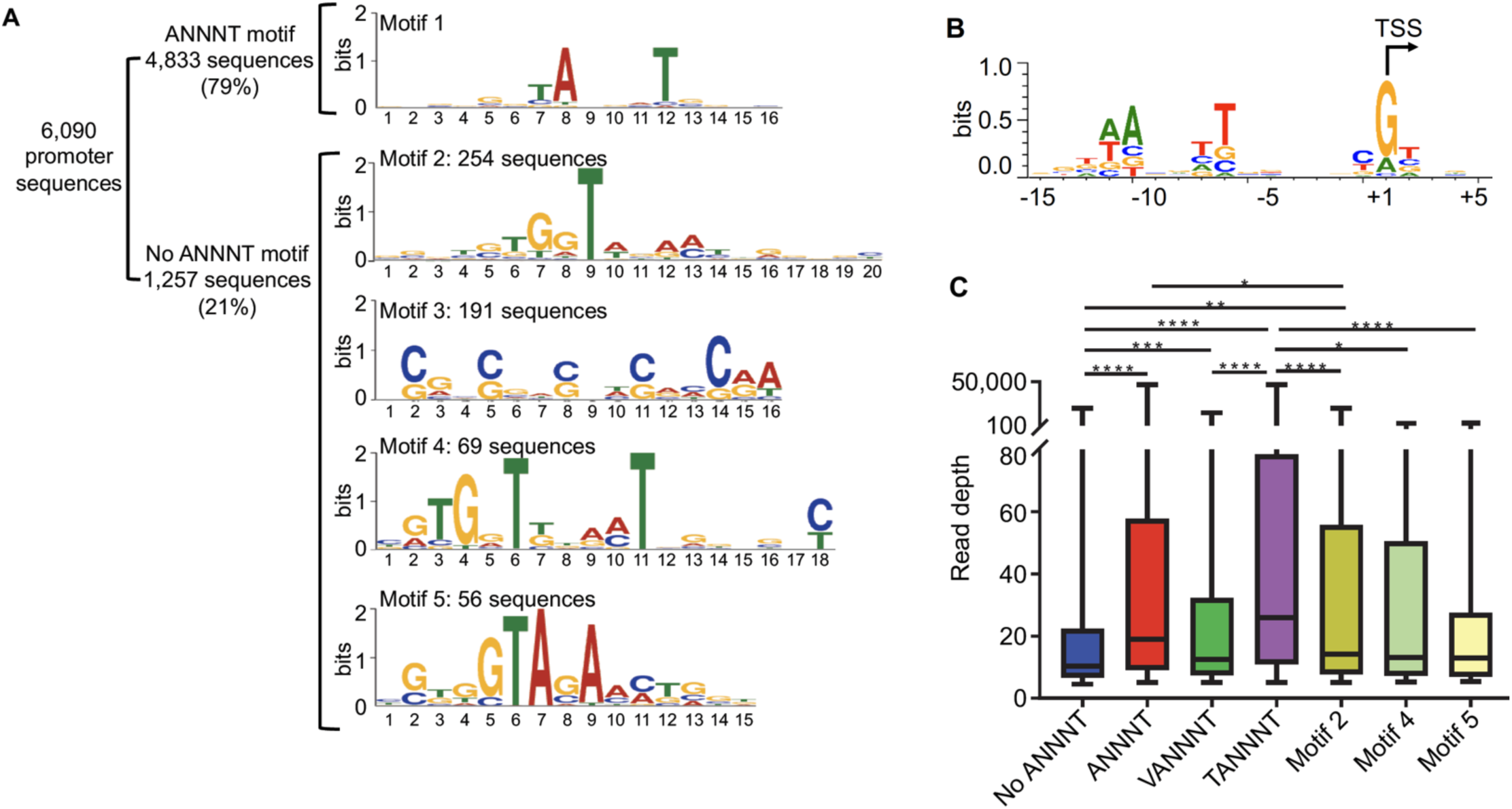
*M. smegmatis* promoter -10 regions are dominated by the ANNNT motif. **A)** Identification of promoter motifs. Consensus motifs were identified by using MEME. The 20 nt upstream the 6,090 TSSs were used for the initial analysis. Those sequences lacking an ANNNT -10 motif between positions -13 and -6 (1,257) were used to identify other conserved promoter sequences. Motif 2 (20 nt length) and Motif 4 (18 nt length) are located immediately upstream of the TSS (at the -1 position), while the spacing of Motif 5 varies from -4 to -1 relative to the TSS, with -3 being the dominant position (75% of the motifs). **B)** The sequences flanking 3,500 randomly chosen TSSs were used to create a sequence logo by WebLogo 3 (Crooks et al, 2004), revealing the two dominant spacings for the ANNNT motif and base preferences in the immediate vicinity of the TSS. **C)** Comparison of apparent promoter activity for different motifs. Mean normalized read depth in the converted libraries from Dataset 1 was compared for TSSs having or lacking the ANNNT motif in the -10 region, and ANNNT-associated TSSs were further subdivided into those containing the extended TANNNT motif or conversely the VANNNT sequence (where V = A, G or C). Motifs 2, 4 and 5 in Figure 2A are also included. ^⋆⋆⋆⋆^p <0.0001, ^⋆⋆⋆^p <0.001, ^⋆⋆^p <0.01, ^⋆^p <0.05 (Kruskal-Wallis test with post-test for multiple comparisons).

Interestingly, we identified several alternative motifs in the -10 promoter regions of transcripts lacking the ANNNT motif (**Figure 2A**). One of these, (G/C)NN(G/C)NN(G/C), is likely the signature of *M. smegmatis’* codon bias in the regions upstream of iTSSs. The other three sequences are candidate binding sites for alternative sigma factors, which are known to be important in regulation of transcription under diverse environmental conditions. However, the identified consensus sequences differ substantially from those previously described in mycobacteria (Raman *et al.*, 2001, Raman *et al.*, 2004, Sun *et al.*, 2004, Lee *et al.*, 2008, Lee *et al.*, 2008, Song *et al.*, 2008, Veyrier *et al.*, 2008, Humpel *et al.*, 2010, Gaudion *et al.*, 2013). The TSSs having these sigma factor motifs and the associated genes are listed in **Table S6.** We next examined the relationship between promoter sequence and promoter strength, as estimated by the read depths in the 5’ end converted libraries. As shown in **Figure 2C**, the expression levels of transcripts with ANNNT -10 motifs are on average substantially higher than those lacking this sequence. In addition, promoters with the full TANNNT motif are associated with more highly abundant transcripts compared to those having a VANNNT sequence, where V is G, A or C. These results implicate TANNNT as the preferred -10 sequence for the housekeeping sigma factor, SigA, in *M. smegmatis*. As shown in Figure 2C, expression levels of transcripts having the motif 2 in **Figure 2A** were significantly increased when compared to those lacking the ANNNT motif.

### 4. Leaderless transcription is a prominent feature of the *M. smegmatis* transcriptome

5’ UTRs play important roles in post-transcriptional regulation and translation, as they may contain regulatory sequences that can affect mRNA stability and/or translation efficiency. Whereas in most bacteria 5’-UTR bearing (“leadered”) transcripts predominate, this is not the case for Mtb, in which near one quarter of the transcripts have been reported to be leaderless (Cortes *et al.*, 2013, Shell *et al.*, 2015). To investigate this feature in *M. smegmatis*, we analyzed the 5’ UTR lengths of all genes that had at least one pTSS. We found that for 24% of the transcripts the TSS coincides with the translation start site or produces a leader length ≤5 nt, resulting in leaderless transcripts (**Figure 3A**). A total of 1,099 genes (including those re-annotated in section 1) have leaderless transcripts, and 155 of those (14%) are also transcribed as leadered mRNAs from separate promoters. For leadered transcripts, the median 5’ UTR length was 69 nt. Interestingly, 15% of the leaders are > 200 nt, suggesting that these sequences may contain potential regulatory elements. We then sought to compare the leader lengths of *M. smegmatis* genes with the leader lengths of their homologs in Mtb. For this analysis we used two independent pTSS mapping Mtb datasets obtained from Cortes et al, 2013 and Shell et al, 2015 (**Figure 3B**). To avoid ambiguities, we used only genes that had a single pTSS in both species. Our results show a statistically significant correlation of leader lengths between species, suggesting that similar genes conserve their transcript features and consequently may have similar regulatory mechanisms. Additionally, comparison of leaderless transcription in *M. smegmatis* and Mtb revealed that 62% or 73% of the genes that are only transcribed as leaderless in *M. smegmatis* also lack a 5’ UTR in MTB, according to Cortes et al, 2013 or Shell et al, 2015, respectively (**Table S7**). We next assessed if leaderless transcripts are associated with particular gene categories, and found the distribution across categories was uneven (**Figure 3C**). The three categories “DNA metabolism,” “Amino acid biosynthesis,” and “Biosynthesis of cofactors, prosthetic groups and carriers” were significantly enriched in leaderless transcripts (*p*-value < 0.05, hypergeometric test), while “Signal transduction,” “Transcription,” and “Transport and binding proteins” appear to have less leaderless transcripts.

**Figure 3.**
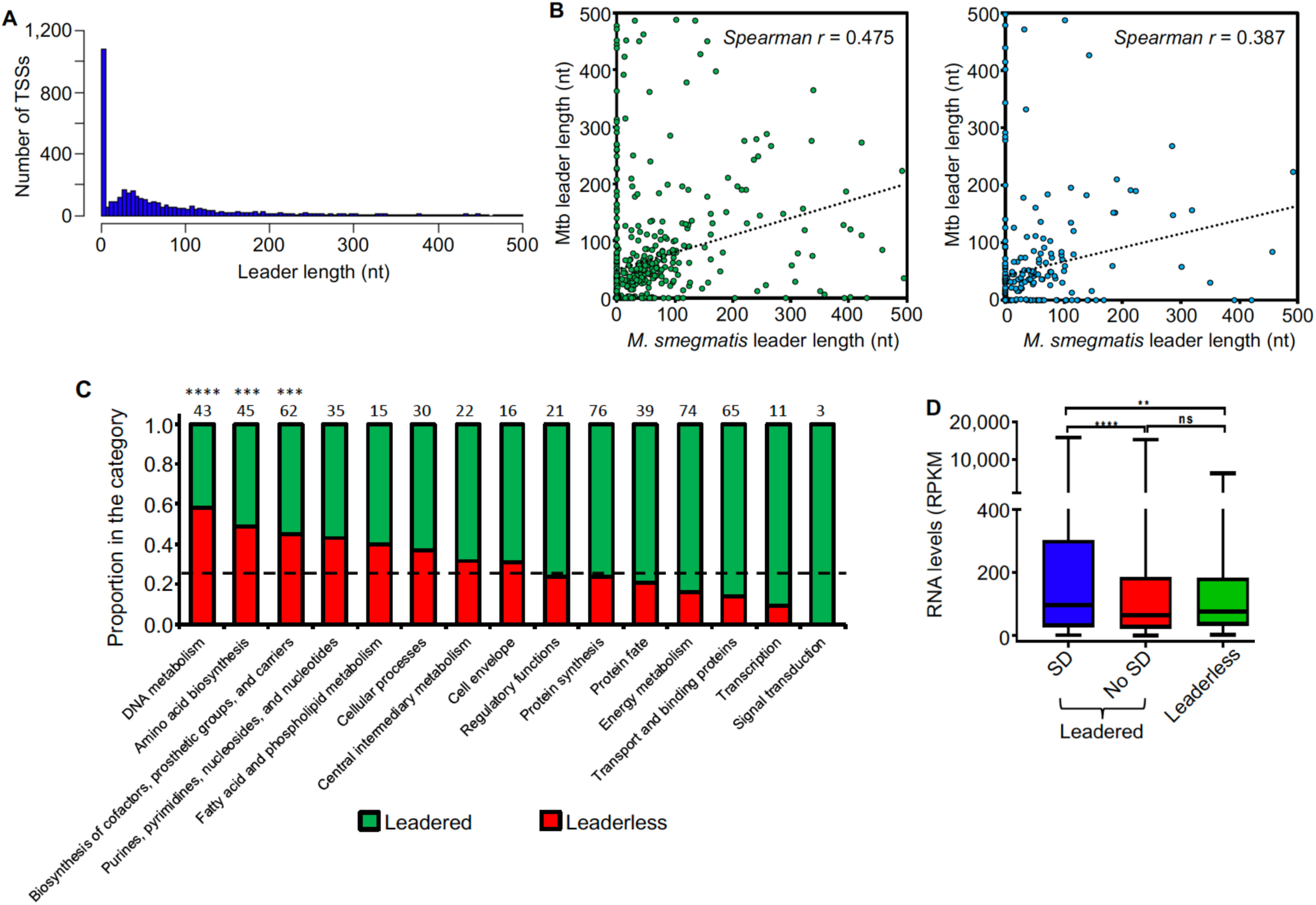
Leader features are conserved in mycobacteria. **A)** Leader length distribution. The 4,054 pTSSs and the pTSSs of the 213 reannotated genes (N-iTSSs→pTSSs) were used for this analysis. **B)** Leader length correlation between *M. smegmatis* and Mtb genes. The leader sequences of genes having a single unique pTSS in both species (leader length ≥0 and ≤500 nt) were used for this analysis. 508 homologous genes in Cortes et al, 2013 (left figure) and 251 homologous genes in Shell et al, 2015 (right figure) were used. When a gene in *M. smegmatis* had more than one homolog in Mtb, that with the highest identity was considered. Spearman r *p*-value <0.00001 in both cases. **C)** Distribution of leaderless transcripts among different functional TIGRfam functional categories. A total of 557 genes having a TIGRfam category were used for this analysis. Genes having both leadered and leaderless transcripts were excluded from this analysis. The black dashed line indicates the expected proportion of leaderless genes (25%) according to the global analysis performed in this study. The numbers above each bar indicate the total number of genes used for this analysis in each category (leaderless + leadered). ^⋆⋆⋆⋆^p <0.0001, ^⋆⋆⋆^p <0.001 (Chi-Square test with Bonferroni correction for multiple comparisons). **D)** RNA levels vary according to leader status. Mean expression levels were compared for genes expressed with leaders containing a canonical SD sequence (SD) or not (No SD) or lacking leaders (leaderless). Gene expression was quantified as RPKMs using RNAseq expression data. Genes were classified as containing an SD sequence if at least one of the three tetramers AGGA, GGAG or GAGG (core sequence AGGAGG) were present in the region -6 to -17 nt relative to the start codon. rRNAs, tRNAs, sRNAs, and genes expressed as both leadered and leaderless transcripts were excluded from this analysis. ^⋆⋆⋆⋆^p <0.0001; ^⋆⋆^p <0.005; ns: not significant. (Kruskal-Wallis test with post-test for multiple comparisons).

We next evaluated the presence of the Shine-Dalgarno ribosome-binding site (SD) upstream of leadered coding sequences. For this analysis, we considered those leaders containing at least one of the three tetramers AGGA, GGAG or GAGG (core sequence AGGAGG) in the region -6 to -17 relative to the start codon to possess canonical SD motifs. We found that only 47% of leadered coding sequences had these canonical SD sequences. Thus, considering also the leaderless RNAs, a large number of transcripts lack canonical SD sequences, suggesting that translation initiation can occur through multiple mechanisms in *M. smegmatis*. We further compared the relative expression levels of leaderless and leadered coding sequences subdivided by SD status. Genes expressed as both leadered and leaderless transcripts were excluded from this analysis. We found that on average, expression levels were significantly higher for those genes with canonical SD sequences than for those with leaders but lacking this motif and for those that were leaderless (**Figure 3D**). Together, these data suggest that genes that are more efficiently translated have also higher transcript levels. Similar findings were made in Mtb, where proteomic analyses showed increased protein levels for genes with SD sequences compared to those lacking this motif (Cortes *et al.*, 2013).

### 5. Identification of novel leaderless ORFs in the *M. smegmatis* genome

As GTG or ATG codons are sufficient to initiate leaderless translation in mycobacteria (Shell *et al.*, 2015, Potgieter *et al.*, 2016), we used this feature to look for unannotated ORFs in the *M. smegmatis* NC_008596 reference genome. Using 1,579 TSSs that remained after pTSS assignment and gene reannotation using N-iTSSs (see **Figure S5**) we identified a total of 66 leaderless ORFs encoding putative proteins longer than 30 amino acids, 5 of which were previously identified (Shell *et al.*, 2015). 83% of these ORFs were predicted in other annotations of the *M. smegmatis* mc^2^155 or MKD8 genome (NC_018289.1, (Gray *et al.*, 2013)), while 10 of the remaining ORFs showed homology to genes annotated in other mycobacterial species and *Helobdella robusta* and two ORFs did not show homology to any known protein. These results show that automatic annotation of genomes can be incomplete and highlight the utility of transcriptomic analysis for genome (re)annotation. Detailed information on these novel putative ORFs is provided in **Table S8**.

### 6. Endonucleolytic RNA cleavage occurs at a distinct sequence motif and is common in mRNA regulatory regions

As our methodology allows us to precisely map RNA cleavage sites in addition to TSSs, we sought to analyze the presence and distribution of cleavage sites in the *M. smegmatis* transcriptome. mRNA processing plays a crucial role in regulation of gene expression, as it is involved in mRNA maturation, stability and degradation (Arraiano *et al.*, 2010). Mixture modeling identified 3,344 CSs with a posterior probability ≥0.9 (high confidence CSs) (**Figure 1A, Table S9**). To determine the sequence context of the CSs, we used the regions flanking the 5’ ends to generate a sequence logo (**Figure 4A**). There was a strong preference for a cytosine in the +1 position (present in more than the 90% of the CSs) (**Figure 4B**), suggesting that it may be structurally important for RNase recognition and/or catalysis.

**Figure 4.**
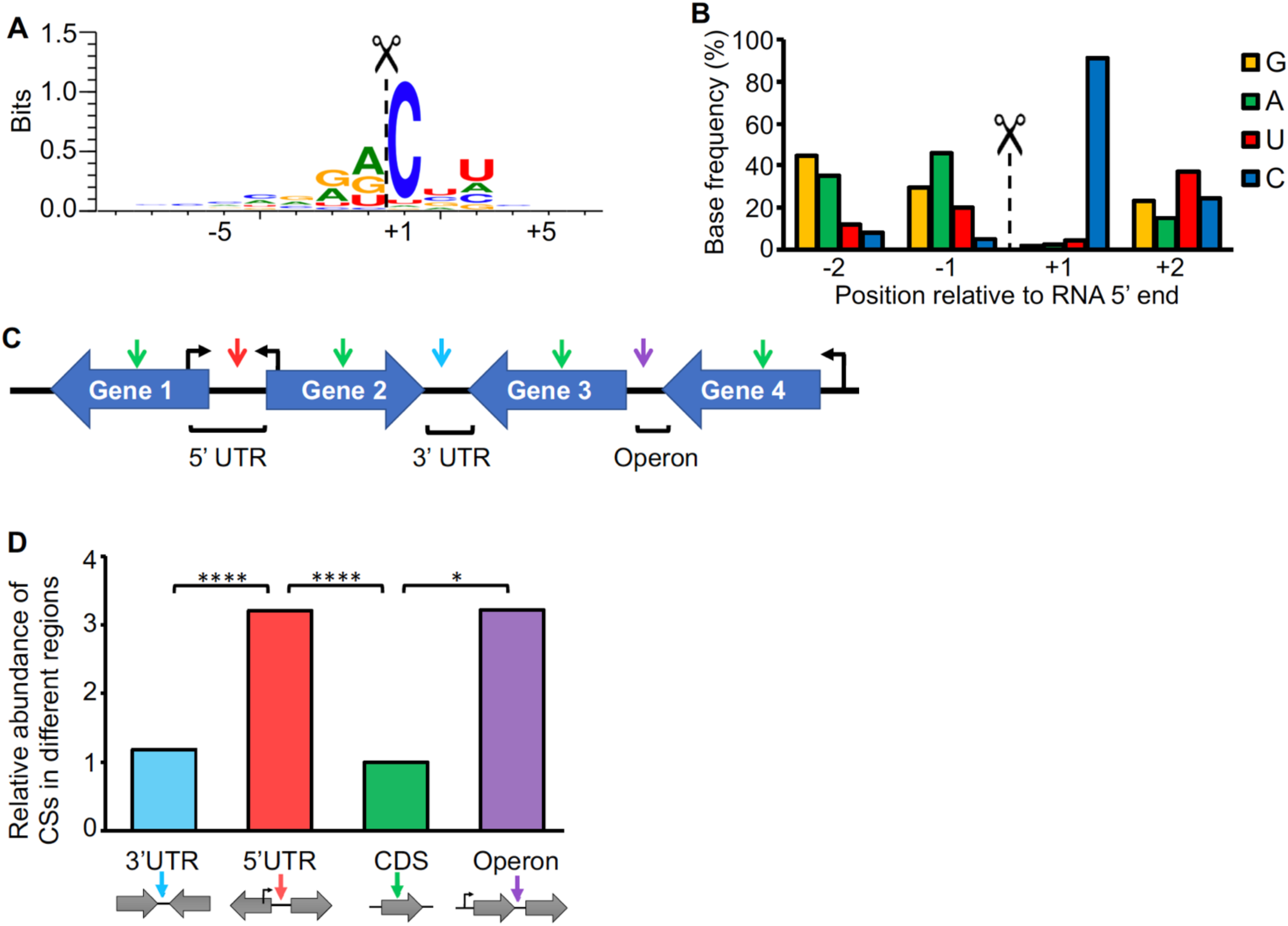
Cleavage site positions are biased with respect to sequence context and genetic location. **A)** Sequence context of cleavage sites. The sequences flanking the 3,344 high-confidence CSs were used to create the sequence logo with WebLogo 3 (Crooks et al, 2004). **B)** Base preference for RNA cleavage. The base frequencies for the -2 to +2 positions were determined. **C)** Cleavage site categories based on the genetic context. CSs are denoted with arrows. 5’ UTR: the CS is within the leader of a gene, and the genes upstream and downstream of the CS are divergent (Gene 1 and Gene 2, red arrow). CDS: The CS is within a coding sequence (green arrow). 3’ UTR: the genes upstream and downstream of the CS are convergent (Gene 2 and Gene 3, light blue arrow). Operon: The CS is between two genes with the same orientation and the first gene in the operon has a pTSS according to Table S5 (violet arrow). **D)** Distribution of cleavage sites. The frequency of CSs in each location was normalized to the proportion of the genome that the location category comprised. The proportions were then normalized to the CDS category, which was set as 1. ^⋆⋆⋆⋆^p <0001, ^⋆^p <0.01 (Chi-square test).

Cleaved 5’ ends can represent either degradation intermediates or transcripts that undergo functional processing/maturation. In an attempt to investigate CS function, we classified them according to their locations within mRNA transcripts (**Figure 4C, Table S9**). We found that, after normalizing to the proportion of the expressed transcriptome that is comprised by each location category, cleaved 5’ ends are more abundant within 5’ UTRs and intergenic regions of operons than within coding sequences and 3’ UTRs (**Figure 4D**). Stringent criteria were used in these analyses to avoid undesired bias (**Figure 4C** and Materials and Methods). While one would expect the CSs associated with mRNA turnover to be evenly distributed throughout the transcript, enrichment of CSs within the 5’ UTRs as well as between two co-transcribed genes may be indicative of cleavages associated with processing and maturation. Alternatively, these regions may be more susceptible to RNases due to lack of associated ribosomes. Here we predicted with high confidence that at least 101 genes have one or more CSs in their 5’ UTRs **(Table S10)**.

We detected cleaved 5’ ends within the coding sequences of 18% of *M. smegmatis* genes, ranging from 1 to over 40 sites per gene. We analyzed the distribution of CSs within coding sequences (**Figure S7**), taking into consideration the genomic context of the genes. When analyzing the distribution of CSs within the coding sequences of genes whose downstream gene has the same orientation, we observed an increase in CS frequency in the region near the stop codon (**Figure S7.A**). However, when only coding sequences having a downstream gene on the opposite strand (convergent) were considered, the distribution of CSs through the coding sequences was significantly different (*p*-value <0.0001, Kolmogorov-Smirnov D test) with the CSs more evenly distributed throughout the coding sequence (**Figure S7.B**). This suggests that the cleavage bias towards the end of the genes observed in **Figure S7.A** may be due to the fact that many of these CSs are actually occurring in the 5’ UTRs of the downstream genes. In cases where the TSS of a given gene occurs within the coding sequence of the preceding gene, a CS may map to both the coding sequence of the upstream gene and the 5’ UTR of the downstream gene. In these cases, we cannot determine in which of the two transcripts the cleavage occurred. However, cleavages may also occur in polycistronic transcripts. We therefore assessed the distributions of CSs in the operons predicted above. The distribution of CSs in genes co-transcribed with a downstream gene showed a slight increase towards the last part of the gene (**Figure S7.C**). This may reflect cases in which polycistronic transcripts are cleaved near the 3’ end of an upstream gene, as has been reported for the *furA*-*katG* operon, in which a cleavage near the stop codon of *furA* was described (Milano *et al.*, 2001, Sala *et al.*, 2008, Taverniti *et al.*, 2011). The *furA*-*katG* cleavage was identified in our dataset, located 1 nt downstream of the previously reported position. A similar enrichment of CSs towards stop codons was also observed in a recent genome-wide RNA cleavage analysis in *Salmonella enterica* (Chao *et al.*, 2017), although in this case the high frequency of cleavage may be also attributed to the U preference of RNase E in this organism, which is highly abundant in these regions.

### 7. Prediction of additional TSSs and CSs based on sequence context

The sequence contexts of TSSs (**Figure 2B**) and CSs (**Figure 4A**) were markedly different, as G and A were highly preferred in the TSS +1 position whereas C was highly preferred in the CS +1 position, and TSSs were associated with a strong overrepresentation of ANNNT -10 sites while CSs were not. These sequence-context differences not only provide validation of our methodology for distinguishing TSSs from CSs, as discussed above, but also provide a means for making improved predictions of the nature of 5’ ends that could not be categorized with high confidence based on their converted/non-converted library coverage alone. Taking advantage of these differences, we sought to obtain a list of additional putative TSSs and CSs. Thus, of the 5’ ends that were not classified by high confidence by mixture modeling, we selected those that had an appropriately positioned ANNNT motif upstream and a G or an A in the +1 position and classified them as TSSs with medium confidence (**Table S11**). In the same way, 5’ ends with a C in the +1 position and lacking the ANNNT motif in the region upstream were designated as medium confidence CSs (**Table S12**). In this way, we were able to obtain 576 and 4,838 medium confidence TSSs and CSs, respectively. Although we are aware of the limitations of these predictions, these lists of medium confidence 5’ ends provide a resource that may be useful for guiding further studies. 5’ ends that did not meet the criteria for high or medium confidence TSSs or CSs are reported in **Table S13**.

### 8. The transcriptional landscape changes in response to oxygen limitation

We sought to study the global changes occurring at the transcriptomic level in oxygen limitation employing the Wayne model (Wayne & Hayes, 1996) with some modifications (see Materials and Methods). Two timepoints were experimentally determined in order to evaluate transcriptomic changes during the transition into hypoxia (**Figure S1**). A different enzyme was used for conversion of 5’ triphosphates to 5’ monophosphates in these 5’-end libraries, and it appeared to be less effective than the enzyme used for the 5’ end libraries in dataset 1. As a consequence, our ability to distinguish TSSs from CSs *de novo* in these datasets was limited. However, we were able to assess changes in abundance of the 5’ ends classified as high-confidence TSSs or CSs in Dataset 1, as well as identify a limited number of additional TSSs and CSs with high confidence (**Figure S4, Table S1**). Corresponding RNAseq expression libraries revealed that, as expected, a large number of genes were up and downregulated in response to oxygen limitation (**Figure S8, Table S14**). We next investigated the transcriptional changes in hypoxia by assessing the relative abundance of TSSs in these conditions. We found 318 high-confidence TSSs whose abundance varied substantially between exponential phase and hypoxia (**Table S15**). A robust correlation was observed between the pTSS peak height in the 5’-end-directed libraries and RNA levels in the expression libraries for hypoxia (**Figure S9**). In an attempt to identify promoter motifs induced in hypoxia, we analyzed the upstream regions of those TSSs whose abundance increased (fold change ≥2, adjusted *p*-value ≤0.05). Interestingly, we detected a conserved GGGTA motif in the -10 region of 56 promoters induced in hypoxia using MEME (**Figure 5A, Table S15**). This motif was reported as the binding site for alternative sigma factor SigF (Rodrigue *et al.*, 2007, Hartkoorn *et al.*, 2010, Humpel *et al.*, 2010). Additionally, the extended -35 and -10 SigF motif was found in 44 of the 56 promoter sequences. (**Figure 5A, Table S15)**. SigF was shown to be induced in hypoxia at the transcript level in Mtb (Iona *et al.*, 2016) and highly induced at the protein level under anaerobic conditions using the Wayne model in *M. bovis* BCG strain (Michele *et al.*, 1999) (Galagan *et al.*, 2013). In *M. smegmatis*, SigF was shown to play a role under oxidative stress, heat shock, low pH and stationary phase (Gebhard *et al.*, 2008, Humpel *et al.*, 2010, Singh *et al.*, 2015) and *sigF* RNA levels were detected in exponential phase at a nearly comparable level to *sigA* (Singh & Singh, 2008). Here, we did not detect significant changes in expression of the *sigF* gene in hypoxia at the transcript level. However, this is consistent with reported data showing that *sigF* transcript levels remain unchanged under stress conditions in *M. smegmatis* (Gebhard *et al.*, 2008), as it was postulated that SigF is post-transcriptionally modulated via an anti-sigma factor rather than through *sigF* transcription activation (Beaucher *et al.*, 2002). We noted that, in the case of TSSs whose abundance was reduced in hypoxia, almost the totality of the promoters contains the -10 ANNNT σ^70^ binding motif. We then examined the presence of SigF motif in the regions upstream of 5’ ends that were not classified as high confidence TSSs. We speculate that 5’ ends associated with this motif may be potential TSSs triggered by hypoxia. We found 96 additional putative TSSs that were (1) overrepresented in hypoxia and (2) associated with appropriately-spaced SigF motifs (**Table S16**). Three of the hypoxia-induced genes with SigF motifs have homologous genes induced in hypoxia in Mtb (Park et al., 2003, Rustad et al., 2008).

**Figure 5.**
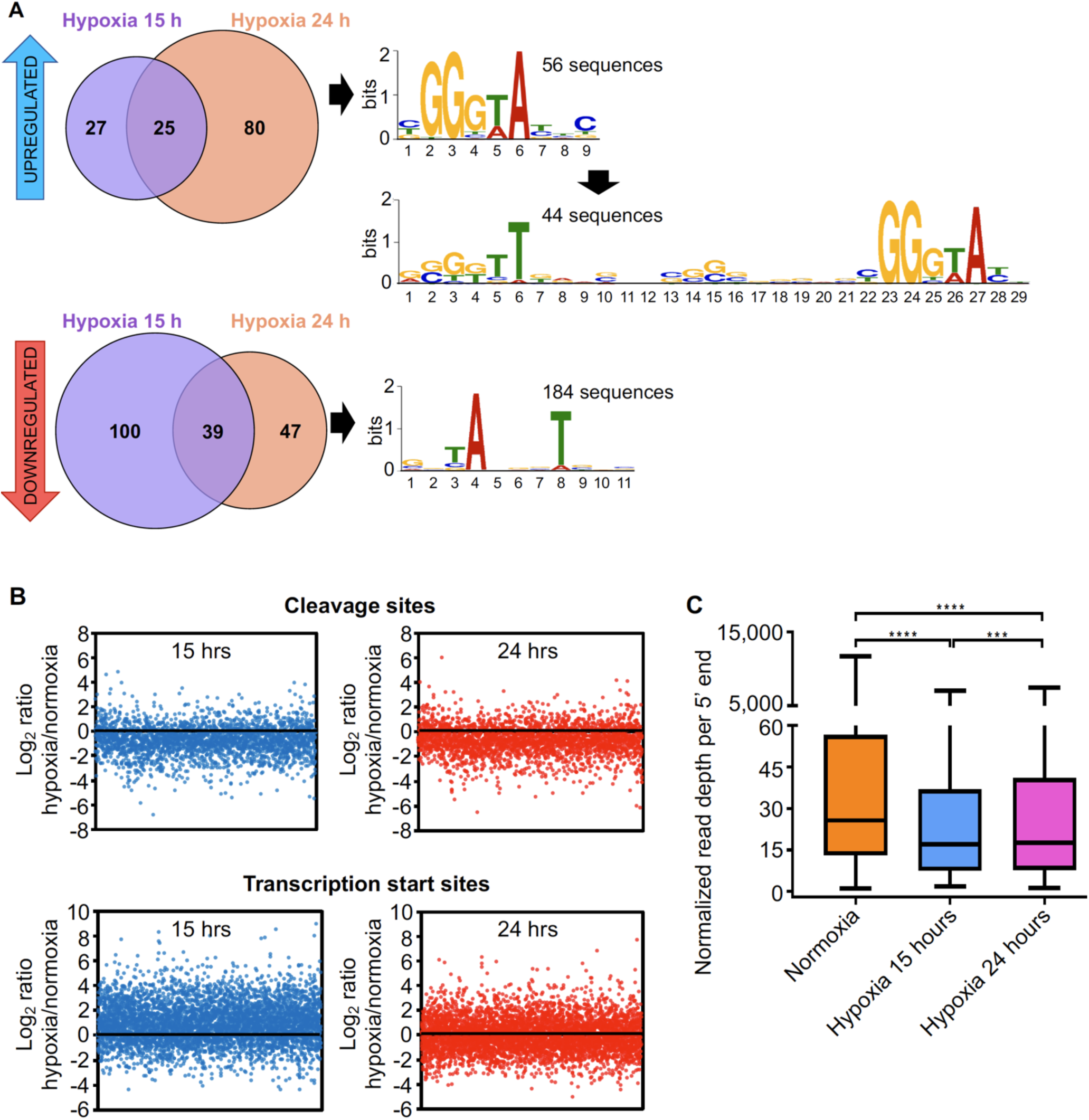
The transcriptional landscape substantially changes upon oxygen limitation. **A)** TSSs significantly increased or decreased in hypoxia. 132 TSSs were overrepresented (upper panel) and 186 were underrepresented (lower panel) in different hypoxia stages. The upstream regions of these TSSs were used to search for promoter motifs using MEME. **B)** The mean normalized read depths for each 5’ end in the non-converted libraries were compared between hypoxia and normoxia. Graphics show the Log2 of the ratios of read depth for each CSs at 15 h (upper left) and 24 h (upper right), and the Log2 of the ratios of the read depth for each TSSs at 15 h (lower left) and 24 h (lower right) compared to normoxia. **C)** Normalized read depth at high-confidence cleavage sites under normoxia and the transition into hypoxia. ^⋆⋆⋆⋆^p <0.0001, ^⋆⋆⋆^p <0.001, ns: not significant (Non-parametric Wilcoxon matched-pairs signed rank test).

It is well known that under anaerobic conditions mycobacteria induce the DosR regulon, a set of genes implicated in stress tolerance (Rosenkrands *et al.*, 2002, O’Toole *et al.*, 2003, Park *et al.*, 2003, Roberts *et al.*, 2004, Rustad *et al.*, 2008, Honaker *et al.*, 2009, Leistikow *et al.*, 2010). The DosR transcriptional regulator was highly upregulated at both hypoxic timepoints in the expression libraries (13 and 18-fold at 15 and 24 hours, respectively). Thus, we hypothesized that the DosR binding motif should be present in a number of regions upstream the TSSs that were upregulated in hypoxia. Analysis of the 200 bp upstream the TSSs using the CentriMo tool for local motif enrichment analysis (Bailey & Machanick, 2012) allowed us to detect putative DosR motifs in 13 or 53 promoters, depending on whether a stringent (GGGACTTNNGNCCCT) or a weak (RRGNCYWNNGNMM) consensus sequence was used as input (Lun *et al.*, 2009, Berney *et al.*, 2014, Gomes *et al.*, 2014) (**Table 15**). At least two of the 13 genes downstream of these TSSs were previously reported to have DosR motifs by Berney and collaborators (Berney *et al.*, 2014) and RegPrecise Database (Novichkov *et al.*, 2013) and two others are homologs of genes in the Mtb DosR regulon that were not previously described in *M. smegmatis* as regulated by DosR (**Table S15**).

We then used CentriMo to search for DosR motifs in the regions upstream of 5’ ends that were not classified as high confidence TSSs, given that TSSs derived from hypoxia-specific promoters may have been absent from Dataset 1. We found 36 putative TSSs associated with 20 different genes (**Table S17**), of which 11 have been shown to have DosR binding motifs (Berney *et al.*, 2014). Five of these are homologs of genes in the Mtb DosR regulon.

#### 9. *M. smegmatis* decreases RNA cleavage under oxygen limitation

There is evidence that mRNA is broadly stabilized under hypoxia and other stress conditions (Rustad *et al.*, 2013, Ignatov *et al.*, 2015). Thus, we anticipated that RNA cleavage should be reduced under hypoxia as a strategy to stabilize transcripts. We compared the relative abundance of each high confidence CS in stress and in exponential phase (**Figure 5B**) and found that RNA cleavage is significantly reduced in both hypoxia 15h and 24h on a global scale (**Figure 5C**). In contrast, relative abundance of TSSs did not decrease in these conditions, indicating that the reduction in CSs is not an artefact of improper normalization (**Figure 5B**). When the ratios of CSs abundance in hypoxia/normal growth of individual genes were analyzed, we observed the same behavior (**Figure S10**). These results indicate that the number of cleavage events per gene decreases during adaptation to hypoxia, which could contribute to the reported increases in half-life (Rustad et al, 2012).

### Discussion

In the past years, genome-wide transcriptome studies have been widely used to elucidate the genome architecture and modulation of transcription in different bacterial species (Albrecht *et al.*, 2009, Mendoza-Vargas *et al.*, 2009, Mitschke *et al.*, 2011, Cortes *et al.*, 2013, Schlüter *et al.*, 2013, Dinan *et al.*, 2014, Ramachandran *et al.*, 2014, Innocenti *et al.*, 2015, Sass *et al.*, 2015, Thomason *et al.*, 2015, Berger *et al.*, 2016, Čuklina *et al.*, 2016, D’arrigo *et al.*, 2016, Heidrich *et al.*, 2017, Li *et al.*, 2017, Zhukova *et al.*, 2017). Here we combined 5’-end-directed libraries and RNAseq expression libraries to shed light on the transcriptional and post-transcriptional landscape of *M. smegmatis* in different physiological conditions.

The implementation of two differentially treated 5’-end libraries followed by Gaussian mixture modeling analysis allowed us to simultaneously map and classify 5’ ends resulting from nucleolytic cleavage and those resulting from primary transcription with high confidence. We were able to classify 57% of the 5’ ends in Dataset 1 with high confidence. In addition, we elaborated a list of medium confidence TSSs and CSs (**Tables S11** and **S12**). These lists constitute a valuable resource for the research community.

Analysis of TSS mapping data allowed us to identify over 4,000 primary TSSs and to study the transcript features in *M. smegmatis*. The high proportion of leaderless transcripts, the lack of a consensus SD sequence in half of the leadered transcripts, and the absence of a conserved -35 consensus sequence indicate that the transcription-translation machineries are relatively robust in *M. smegmatis*. The robustness of transcription and translation are features shared with Mtb, where 25% of the transcripts lack a leader sequence (Cortes *et al.*, 2013, Shell *et al.*, 2015). In addition, high abundances of transcripts lacking 5’ UTRs have been reported in other bacteria including *Corynebacterium diphtheria*, *Leptospira interrogans*, *Borrelia burgdorferi*, and *Deinococcus deserti*, the latter having 60% leaderless transcripts (de Groot *et al.*, 2014, Adams *et al.*, 2017, Zhukova *et al.*, 2017, Wittchen *et al.*, 2018). Considering the high proportion of leaderless transcripts and the large number of leadered transcripts that lack a SD sequence (53%), it follows that an important number of transcripts are translated without canonical interactions between the mRNA and anti-Shine-Dalgarno sequence, suggesting that *M. smegmatis* has versatile mechanisms to address translation. A computational prediction showed that the presence of SD can be very variable between prokaryotes, ranging from 11% in Mycoplasma to 91% in Firmicutes (Chang *et al.*, 2006). Cortes et al (Cortes *et al.*, 2013) reported that the 55% of the genes transcribed with a 5’ UTR lack the SD motif. These similarities between *M. smegmatis* and *M. tuberculosis*, along with the correlation of leader lengths for homologous genes between species shown in **Figure 3B**, provide further evidence that many transcriptomic features are conserved between mycobacterial genomes. These data support the idea that in many cases, similar mechanisms govern modulation of gene expression in both species.

In an attempt to understand the role of RNA cleavage in mycobacteria, we identified and classified over 3,000 CSs throughout the *M. smegmatis* transcriptome, presenting the first report of an RNA cleavage map in mycobacteria. The most striking feature of the CSs was a cytidine in the +1 position, which was true in over 90% of the cases. While the RNases involved in global RNA decay in mycobacteria have not been yet elucidated, some studies have implicated RNase E as a major player in RNA processing and decay (Kovacs *et al.*, 2005, Zeller *et al.*, 2007, Csanadi *et al.*, 2009, Taverniti *et al.*, 2011), given its central role in other bacteria such as *E. coli* and its essentiality for survival in both *M. smegmatis* and Mtb (Sassetti *et al.*, 2003, Sassetti & Rubin, 2003, Griffin *et al.*, 2011, Taverniti *et al.*, 2011, DeJesus *et al.*, 2017). It is therefore possible that mycobacterial RNase E, or other endonucleases with dominant roles, favor cytidine in the +1 position. Interestingly, the sequence context of cleavage found here is different from that described for *E. coli*, for which the consensus sequence is (A/G)N↓AU (Mackie, 2013) or *S. enterica*, in which a marked preference for uridine at the +2 position and AU-rich sequences are important for RNase E cleavage (Chao *et al.*, 2017).

RNA cleavage is required for maturation of some mRNAs (Li & Deutscher, 1996, Condon *et al.*, 2001, Gutgsell & Jain, 2010, Moores *et al.*, 2017). Therefore, the observation that CSs are enriched in 5’ UTRs and intergenic regions suggests that processing may play roles in RNA maturation, stability, and translation for some transcripts in *M. smegmatis*. A high abundance of processing sites around the translation start site was also observed in *P. aeruginosa* and *S. enterica* in transcriptome-wide studies (Chao *et al.*, 2017, Gill *et al.*, 2018), suggesting that 5’ UTR cleavage may be a widespread post-transcriptional mechanism for modulating gene expression in bacteria.

Regulation of RNA decay and processing plays a crucial role in adaptation to environmental changes. We present evidence showing that RNA cleavage is markedly reduced in conditions that result in growth cessation. It was previously demonstrated that in low oxygen concentrations mycobacteria reduce their RNA levels (Ignatov *et al.*, 2015) and mRNA half-life is strikingly increased (Rustad *et al.*, 2013), likely as a mechanism to maintain adequate transcript levels in the cell without the energy expenditures that continuous transcription would require. While several traits are involved in the regulation of transcript abundance and stability, the observation that cleavage events are pronouncedly reduced in these conditions pinpoint this mechanism as a potential way to control RNA stability under stress. In agreement with this hypothesis, RNase E was modestly but significantly decreased at the transcript level in early and late hypoxia (fold change = 0.63 and 0.56, respectively, *p*-value adjusted <0.05), suggesting that reducing the RNase E abundance in the cell may be a strategy to increase transcript half-life. Further study is needed to better understand the relationship between transcript processing and RNA decay in normoxic growth as well as stress conditions.

Hypoxic stress conditions were also characterized by major changes in the TSSs. 5’-end-mapping libraries revealed that over 300 TSSs varied substantially when cultures were limited in oxygen. We found that 56 transcripts triggered in hypoxia contain the SigF promoter binding motif, indicating that this sigma factor plays a substantial role in the *M. smegmatis* hypoxia response. While previous work revealed increased expression of SigF itself in hypoxia in Mtb (Galagan *et al.*, 2013, Iona *et al.*, 2016, Yang *et al.*, 2018), this is the first report demonstrating the direct impact of SigF on specific promoters in hypoxic conditions in mycobacteria. Further work is needed to better understand the functional consequences of SigF activation in both organisms in response to hypoxia.

## Acknowledgements

This work was supported by NSF CAREER award 1652756 to SSS. We are grateful to Zheyang Wu and Thomas Ioerger for helpful advice on the mixture modeling and Michael Chase for helpful advice on other aspects of our data analysis pipeline. We thank members of the Shell lab for technical assistance and helpful discussions. All Illumina sequencing was performed by the UMass Medical School Deep Sequencing Core.

## Supplementary Figures

**Supplementary Figure 1.**
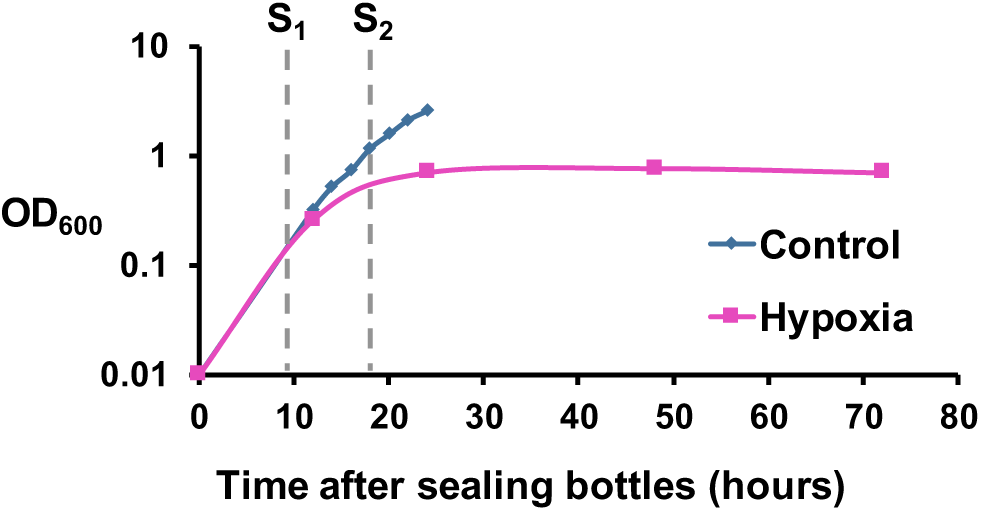
Wayne hypoxia model. Cultures were grown in sealed flasks to produce a gradual reduction in oxygen. Samples were taken at 15 (S1) and 24 (S2) hours after bottles were sealed. For control, cultures were sampled at an OD = 0.8.

**Figure S2.**
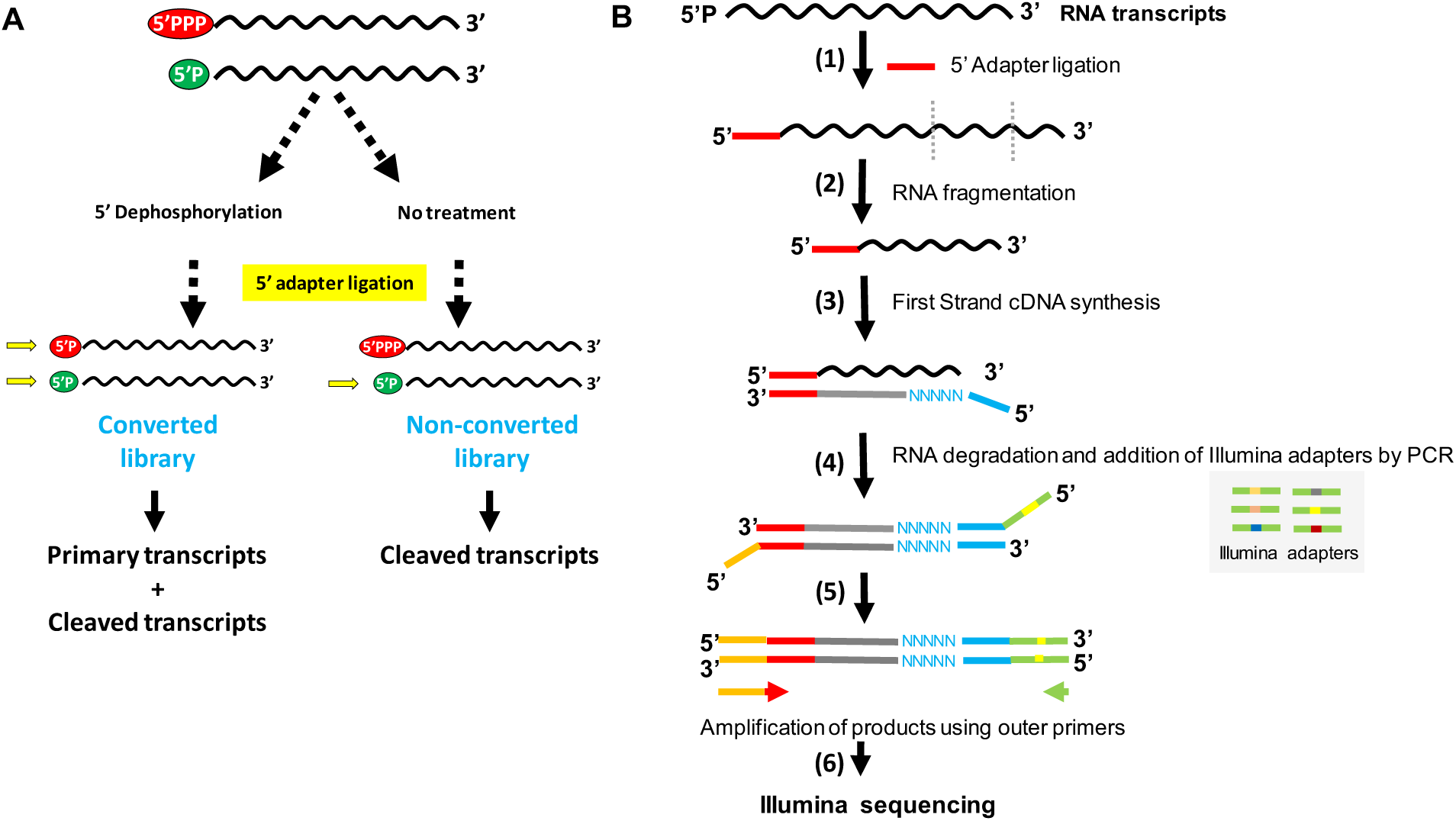
Construction of 5’-end-directed libraries. **A)** RNA samples were split in two parts and treated differentially. RNA for Library 1 (converted) was treated with RPPH to convert triphosphates in monophosphates, allowing the capture of 5’ end that are primary transcripts or cleaved RNAs. RNA for Library 2 (non-converted) was mock-treated, allowing the capture of cleaved transcripts. **B)** Workflow of 5’-end-directed libraries. After RPPH or 5’ polyphosphatase treatment, adapter SSS392 (TCCCTACACGACGCTCTTCCGAUCU) was ligated to the 5’ monophosphate ends (1). Then, RNA was fragmented by heating at 85°C for 6 min (log phase experiment) or at 94°C for 11 min (hypoxia experiment) (2) and first strand cDNA synthesis was carried out using the degenerate primer SSS397 (CTGGAGTTCAGACGTGTGCTCTTCCGATCTNNNNNN) (3). RNA was then degraded and DNA was amplified using universal adapter sequence SSS398 (AATGATACGGCGACCACCGAGATCTACACTCTTTCCCTACACGACGCTCTTC) and primers bearing Illumina indexes (4). Adapter-bearing products were PCR-amplified using outer primers SSS401 (AATGATACGGCGACCACCGAGATC) and SSS402 (CAAGCAGAAGACGGCATACGAGAT) to enrich for full-length fragments. 4 (log phase experiment) or 16 (hypoxia experiment) PCR cycles were performed (5). Finally, libraries were sequenced using Illumina technology (6).

**Figure S3.**
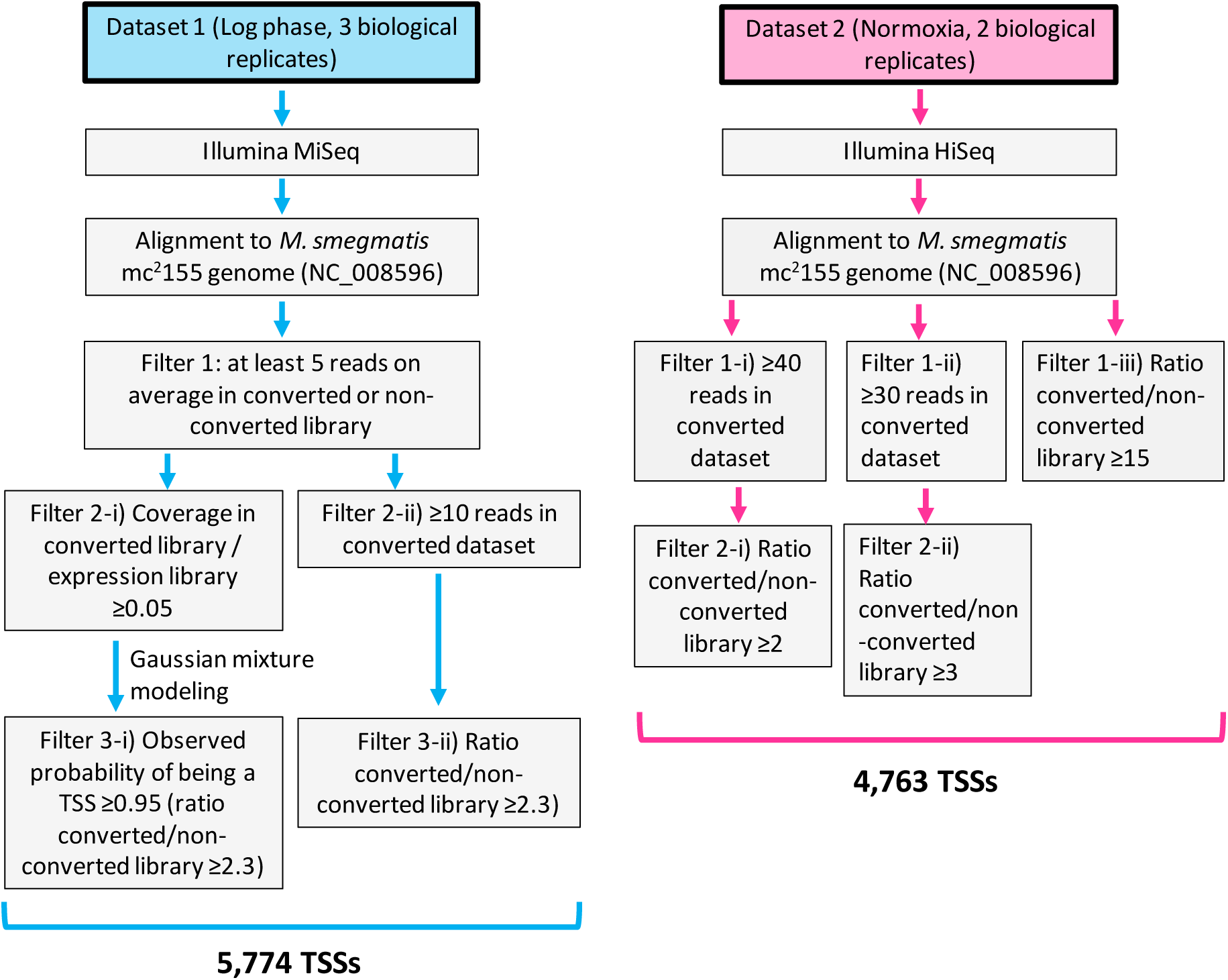
Workflow for noise filtering and TSS prediction in the different datasets.

**Supplementary Figure 4.**
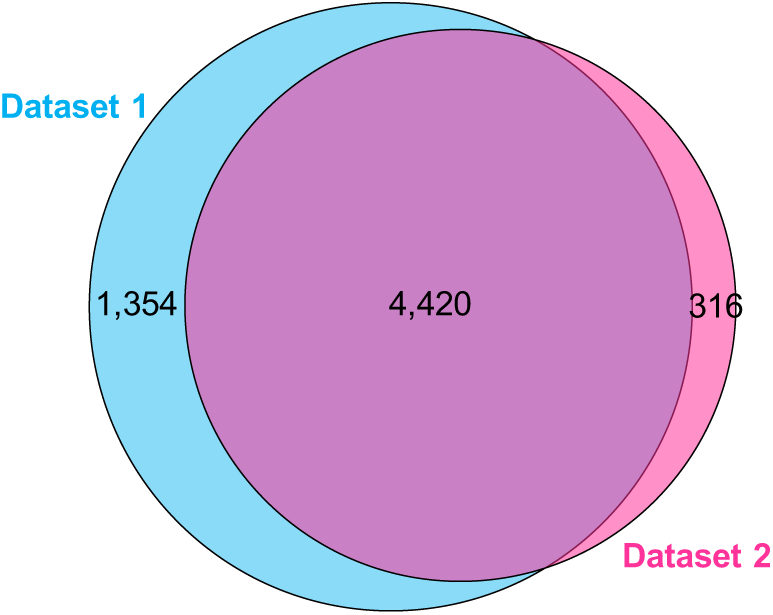
TSSs identified in the different datasets. Dataset 1: exponential phase (5,774 TSSs), Dataset 2: a separate exponential phase dataset (aka normoxia) obtained as a control for a matched hypoxia dataset (4,736 TSSs).

**Figure S5.**
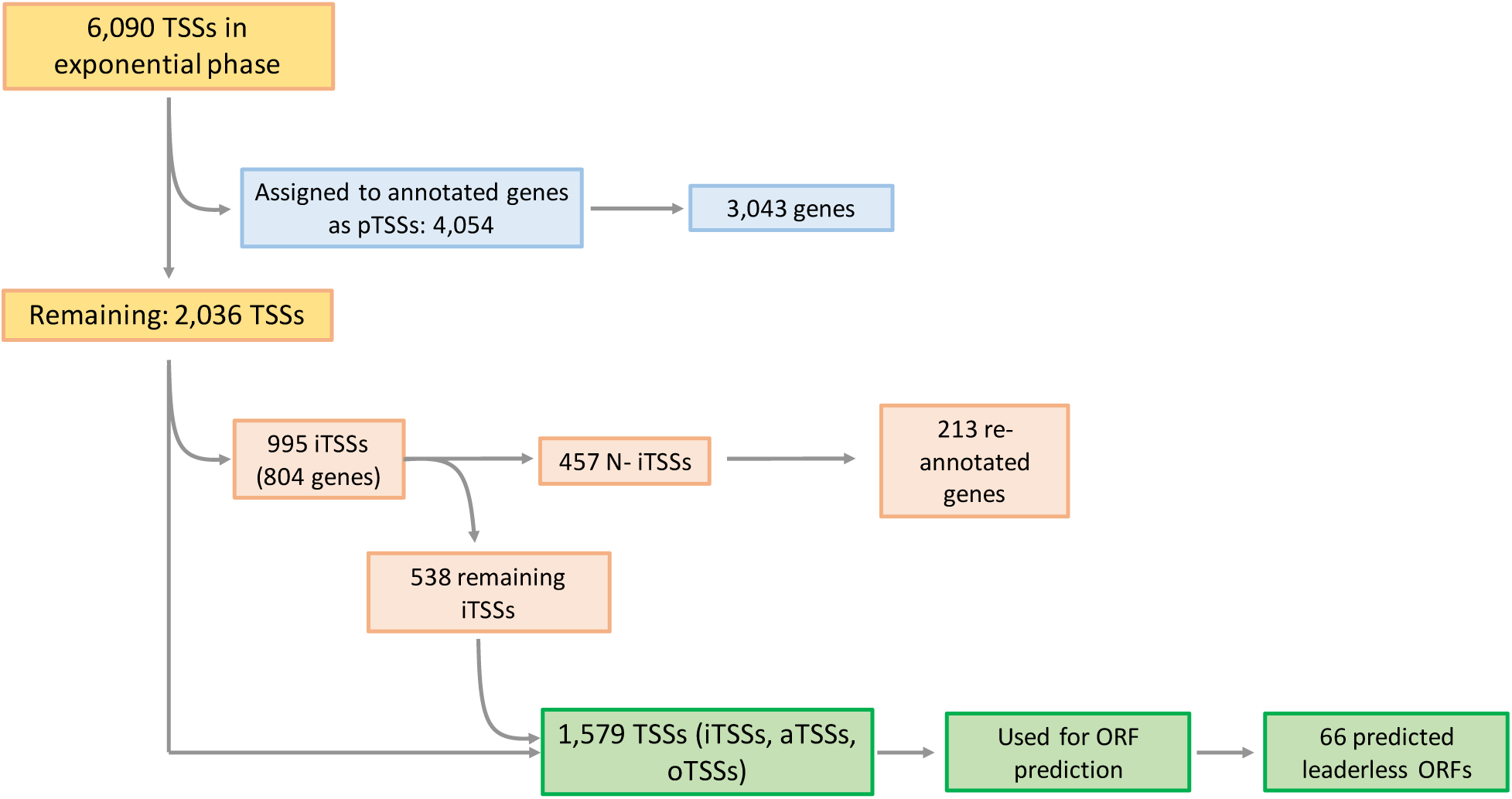
Workflow used for TSS classification. A complete scheme of the procedure used to classify TSSs is shown. TSSs located within 0-500 nt upstream of an annotated coding sequence were classified as pTSSs. TSSs located within annotated coding sequences were classified as iTSSs. iTSSs located within the first 25% of an annotated coding sequence were subclassified as N-iTSSs. When a gene lacked a pTSS, had an N-iTSS, and had an in-frame start codon downstream of the N-iTSS and within the first 30% of the coding sequence, the start codon of the gene was re-annotated. aTSSs (TSSs located on the antisense strand of a coding sequence, 5’ UTR, or 3’ UTR) and oTSSs (TSSs not belonging to any of the above-mentioned categories) were assigned as described in Figure 1D and Materials and Methods.

**Figure S6.**
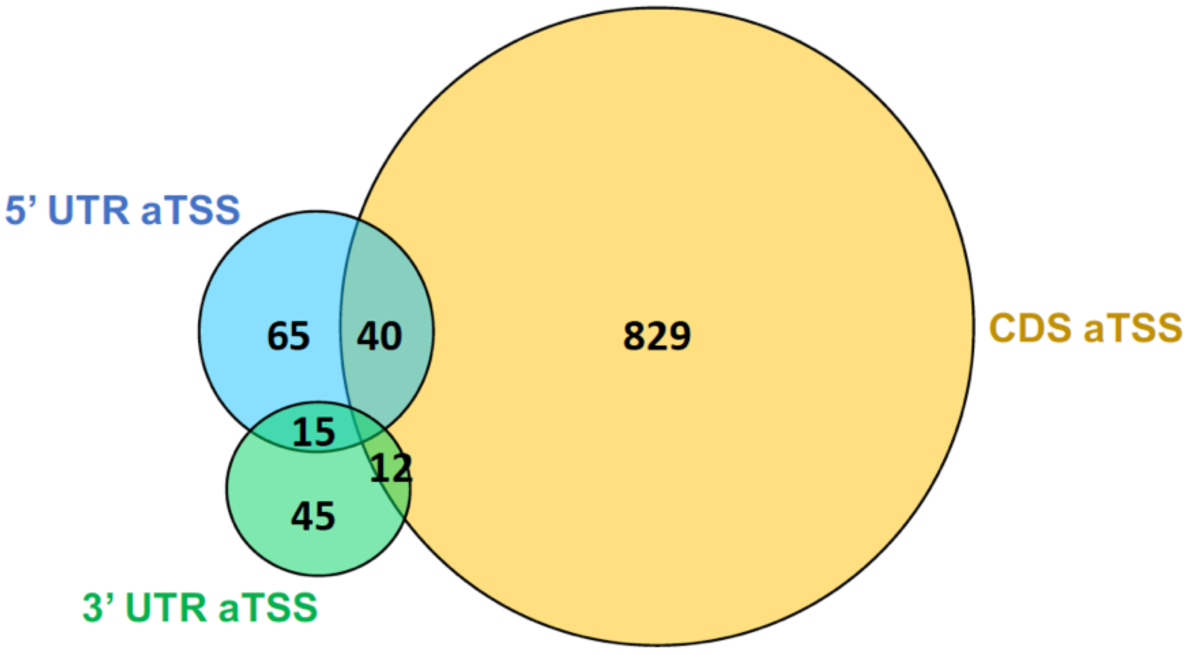
Distribution of antisense TSSs. The 1,006 aTSSs were classified according to their positions in 5’ UTRs, 3’ UTRs, and CDSs (coding sequences).

**Supplementary Figure 7.**
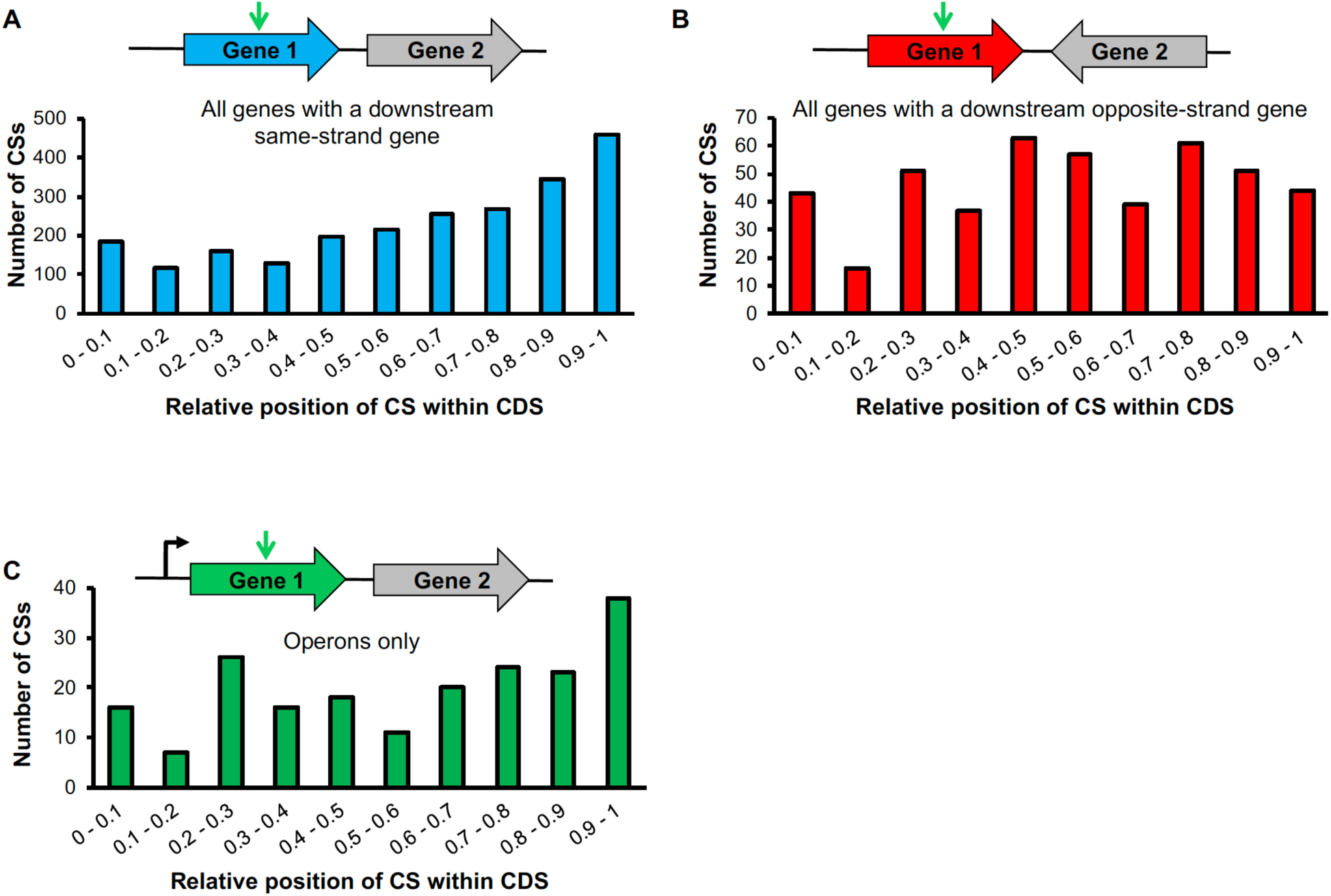
Cleavage sites distribution within genes according to coding sequence context. The number of cleavage sites according to the relative position in the coding sequence is represented considering **A)** only coding sequences whose downstream gene is on the same strand, **B)** only coding sequences whose downstream gene is in the opposite strand (convergent), and **C)** only genes having a downstream gene transcribed as an operon. The CS distribution is significantly different between graphics A and B (*p*-value <0.0001, Kolmogorov Smirnov D test).

**Supplementary Figure 8.**
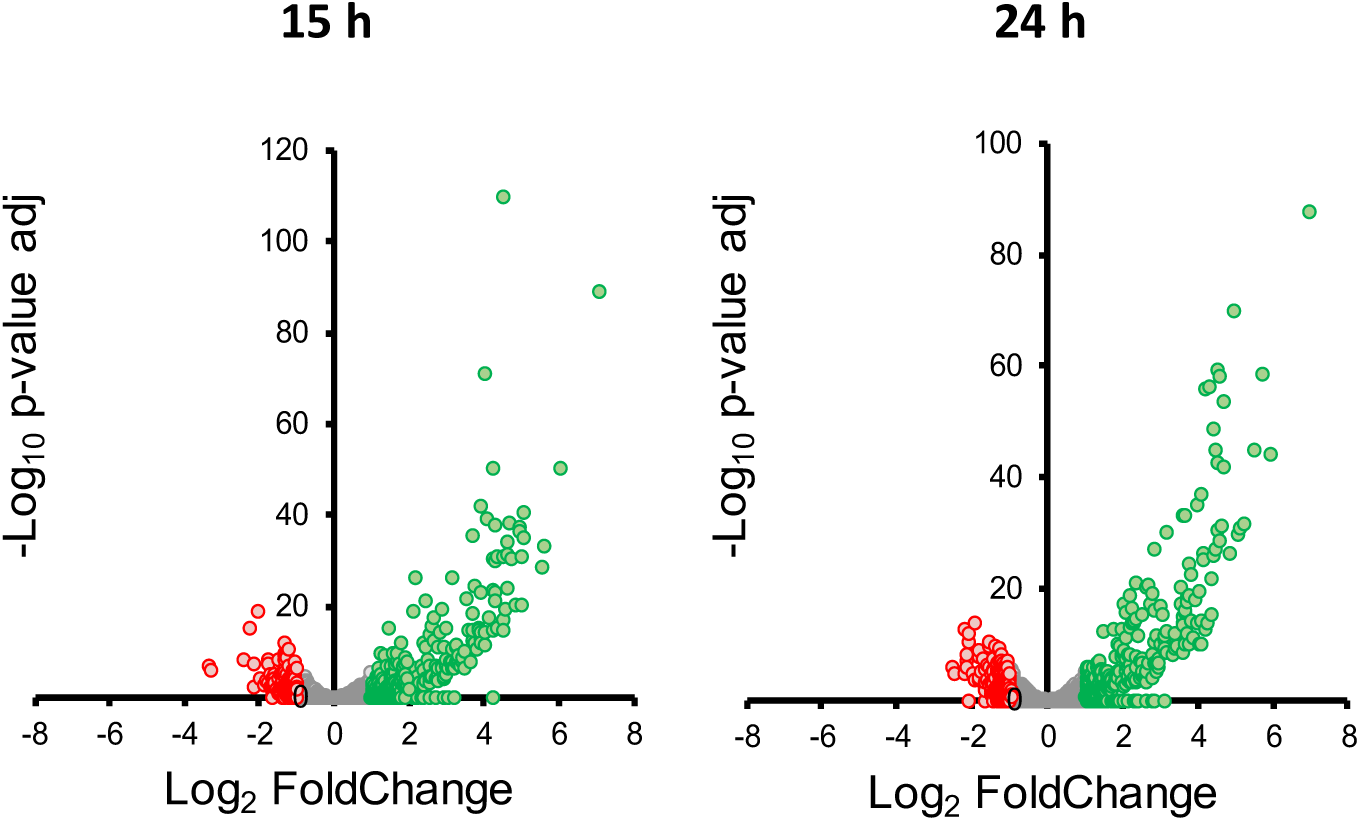
Gene expression levels in RNAseq expression libraries in hypoxia. Changes in transcript levels were obtained by DEseq2 analysis, comparing each indicated condition to the control experiment. Genes upregulated and downregulated with a fold change ≥2 are highlighted in green and red, respectively.

**Supplementary Figure 9.**
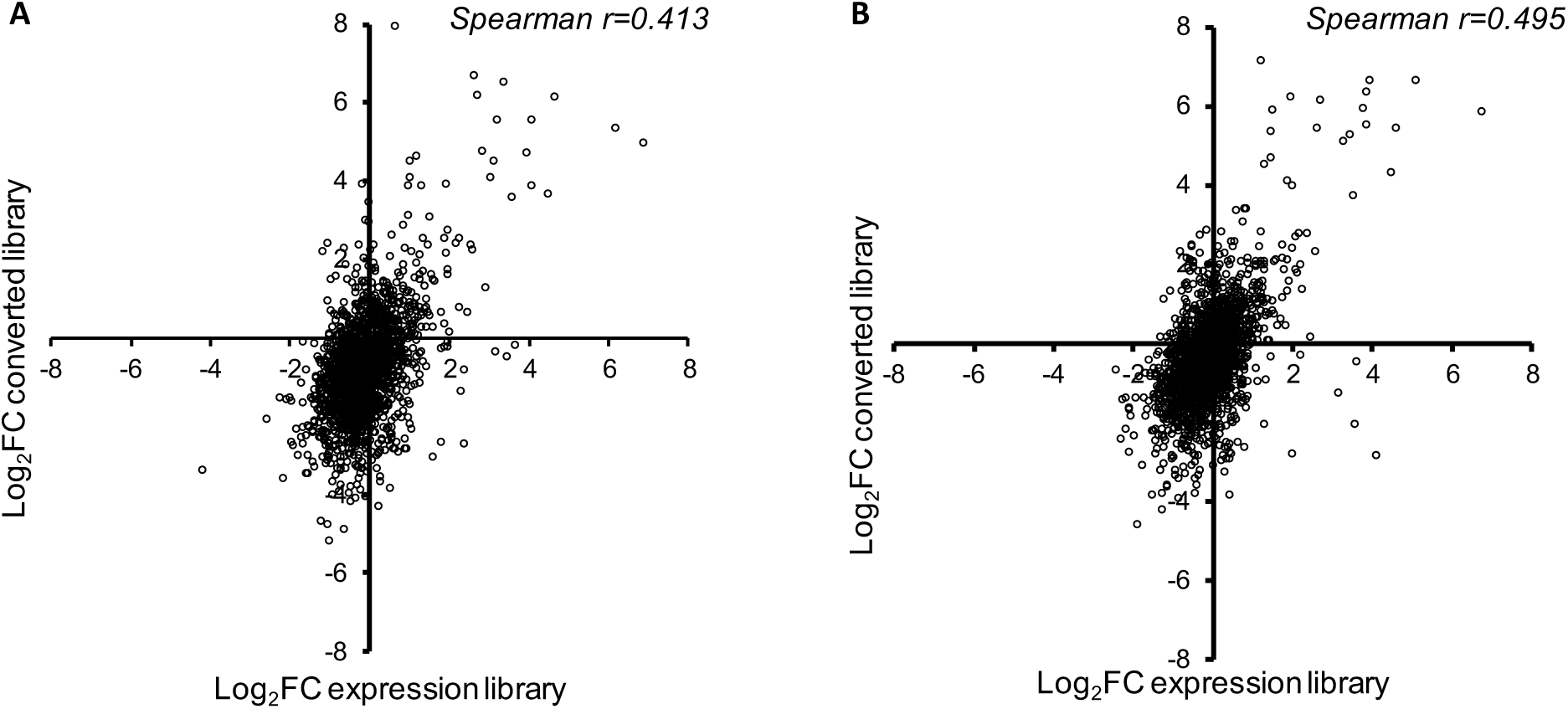
Correlation between expression data and 5’ end-directed libraries data in hypoxia. The *X* axis represents the Log_2_ of the fold change in the expression libraries from hypoxia/normoxia datasets and the *Y* axis represents the Log2 of the fold change in read depth in hypoxia/normoxia 5’-end-directed libraries. The analysis was done for hypoxia at 15 hours **(A)** and 24 hours **(B)**. Genes having only one pTSS were used. The correlation is significant in both cases, with a *p*-value <0.00001.

**Supplementary Figure 10.**
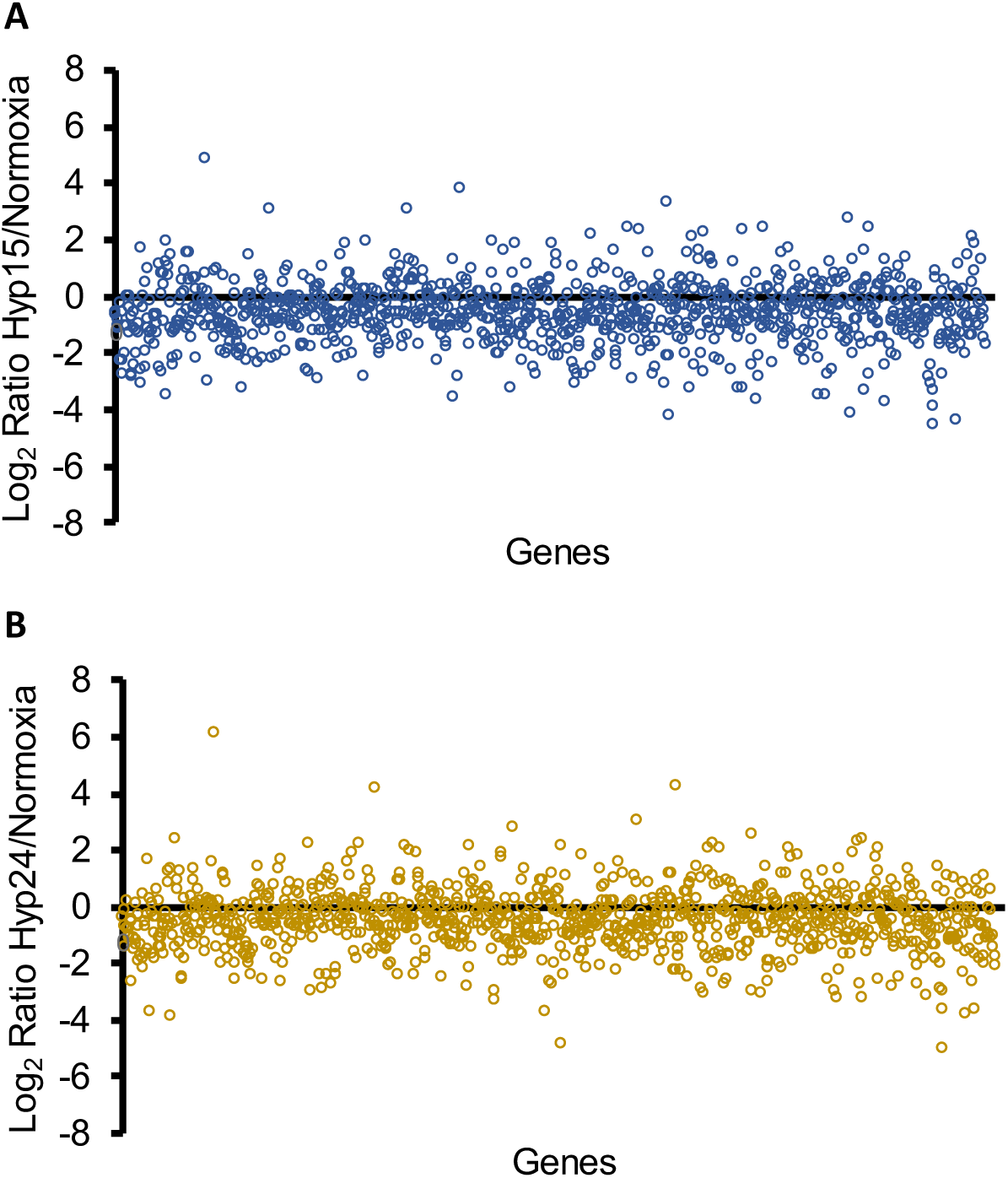
Changes in RNA cleavage within coding sequences in hypoxic conditions. The number of cleavage events within each coding sequence was compared through the different conditions. The Log2 of the ratio of the number of cleavages in hypoxia/control are shown. Each dot represents a specific gene. **A)** Hypoxia 15 hours, **B)** Hypoxia 24 hours.

## References

Adams PP, Flores Avile C, Popitsch N, Bilusic I, Schroeder R, Lybecker M & Jewett MW (2017) In vivo expression technology and 5′ end mapping of the Borrelia burgdorferi transcriptome identify novel RNAs expressed during mammalian infection. Nucleic acids research 45: 775-792.

Albrecht M, Sharma CM, Reinhardt R, Vogel J & Rudel T (2009) Deep sequencing-based discovery of the Chlamydia trachomatis transcriptome. Nucleic acids research 38: 868-877.

Andre G, Even S, Putzer H, Burguiere P, Croux C, Danchin A, Martin-Verstraete I & Soutourina O (2008) Sbox and T-box riboswitches and antisense RNA control a sulfur metabolic operon of Clostridium acetobutylicum. Nucleic Acids Res 36: 5955-5969.

Arraiano CM, Andrade JM, Domingues S, et al. (2010) The critical role of RNA processing and degradation in the control of gene expression. FEMS microbiology reviews 34: 883-923.

Bagchi G, Das TK & Tyagi JS (2002) Molecular analysis of the dormancy response in Mycobacterium smegmatis: expression analysis of genes encoding the DevR–DevS two-component system, Rv3134c and chaperone α-crystallin homologues. FEMS microbiology letters 211: 231-237.

Bailey TL & Machanick P (2012) Inferring direct DNA binding from ChIP-seq. Nucleic Acids Res 40: e128.

Bailey TL, Johnson J, Grant CE & Noble WS (2015) The MEME Suite. Nucleic Acids Res 43: W39-49.

Beaucher J, Rodrigue S, Jacques PE, Smith I, Brzezinski R & Gaudreau L (2002) Novel Mycobacterium tuberculosis anti-sigma factor antagonists control sigmaF activity by distinct mechanisms. Mol Microbiol 45: 1527-1540.

Beaume M, Hernandez D, Farinelli L, Deluen C, Linder P, Gaspin C, Romby P, Schrenzel J & Francois P (2010) Cartography of methicillin-resistant S. aureus transcripts: detection, orientation and temporal expression during growth phase and stress conditions. PLoS One 5: e10725.

Benaglia T, Chauveau D, Hunter DR & Young D (2009) mixtools: An R Package for Analyzing Finite Mixture Models. Journal of Statistical Software 32: 1-29.

Berger P, Knödler M, Förstner KU, Berger M, Bertling C, Sharma CM, Vogel J, Karch H, Dobrindt U & Mellmann A (2016) The primary transcriptome of the Escherichia coli O104: H4 pAA plasmid and novel insights into its virulence gene expression and regulation. Scientific reports 6: 35307.

Berney M, Greening C, Conrad R, Jacobs WR, Jr. & Cook GM (2014) An obligately aerobic soil bacterium activates fermentative hydrogen production to survive reductive stress during hypoxia. Proceedings of the National Academy of Sciences of the United States of America 111: 11479-11484.

Chang B, Halgamuge S & Tang SL (2006) Analysis of SD sequences in completed microbial genomes: non-SD-led genes are as common as SD-led genes. Gene 373: 90-99.

Chao Y, Li L, Girodat D, et al. (2017) In Vivo Cleavage Map Illuminates the Central Role of RNase E in Coding and Non-coding RNA Pathways. Molecular cell 65: 39-51.

Condon C, Brechemier-Baey D, Beltchev B, Grunberg-Manago M & Putzer H (2001) Identification of the gene encoding the 5S ribosomal RNA maturase in Bacillus subtilis: mature 5S rRNA is dispensable for ribosome function. RNA (New York, NY) 7: 242-253.

Coros A, Callahan B, Battaglioli E & Derbyshire KM (2008) The specialized secretory apparatus ESX-1 is essential for DNA transfer in Mycobacterium smegmatis. Molecular microbiology 69: 794-808.

Cortes T, Schubert OT, Rose G, Arnvig KB, Comas I, Aebersold R & Young DB (2013) Genome-wide mapping of transcriptional start sites defines an extensive leaderless transcriptome in Mycobacterium tuberculosis. Cell reports 5: 1121-1131.

Csanadi A, Faludi I & Miczak A (2009) MSMEG_4626 ribonuclease from Mycobacterium smegmatis. Molecular biology reports 36: 2341-2344.

Čuklina J, Hahn J, Imakaev M, Omasits U, Förstner KU, Ljubimov N, Goebel M, Pessi G, Fischer H-M & Ahrens CH (2016) Genome-wide transcription start site mapping of Bradyrhizobium japonicum grown free-living or in symbiosis–a rich resource to identify new transcripts, proteins and to study gene regulation. BMC genomics 17: 302.

D’arrigo I, Bojanovič K, Yang X, Holm Rau M & Long KS (2016) Genome-wide mapping of transcription start sites yields novel insights into the primary transcriptome of Pseudomonas putida. Environmental microbiology 18: 3466-3481.

de Groot A, Roche D, Fernandez B, Ludanyi M, Cruveiller S, Pignol D, Vallenet D, Armengaud J & Blanchard L (2014) RNA sequencing and proteogenomics reveal the importance of leaderless mRNAs in the radiation-tolerant bacterium Deinococcus deserti. Genome biology and evolution 6: 932-948.

DeJesus MA, Gerrick ER, Xu W, et al. (2017) Comprehensive Essentiality Analysis of the Mycobacterium tuberculosis Genome via Saturating Transposon Mutagenesis. MBio 8.

DiChiara JM, Contreras-Martinez LM, Livny J, Smith D, McDonough KA & Belfort M (2010) Multiple small RNAs identified in Mycobacterium bovis BCG are also expressed in Mycobacterium tuberculosis and Mycobacterium smegmatis. Nucleic Acids Res 38: 4067-4078.

Dick T, Lee BH & Murugasu-Oei B (1998) Oxygen depletion induced dormancy in Mycobacterium smegmatis. FEMS microbiology letters 163: 159-164.

Dinan AM, Tong P, Lohan AJ, Conlon KM, Miranda-CasoLuengo AA, Malone KM, Gordon SV & Loftus BJ (2014) Relaxed selection drives a noisy noncoding transcriptome in members of the Mycobacterium tuberculosis complex. MBio 5: e01169-01114.

Elharar Y, Roth Z, Hermelin I, Moon A, Peretz G, Shenkerman Y, Vishkautzan M, Khalaila I & Gur E (2014) Survival of mycobacteria depends on proteasome-mediated amino acid recycling under nutrient limitation. The EMBO journal 33: 1802-1814.

Fields CJ & Switzer RL (2007) Regulation of pyr gene expression in Mycobacterium smegmatis by PyrR-dependent translational repression. J Bacteriol 189: 6236-6245.

Fozo EM, Kawano M, Fontaine F, Kaya Y, Mendieta KS, Jones KL, Ocampo A, Rudd KE & Storz G (2008) Repression of small toxic protein synthesis by the Sib and OhsC small RNAs. Mol Microbiol 70: 1076-1093.

Galagan JE, Minch K, Peterson M, et al. (2013) The Mycobacterium tuberculosis regulatory network and hypoxia. Nature 499: 178-183.

Gaudion A, Dawson L, Davis E & Smollett K (2013) Characterisation of the Mycobacterium tuberculosis alternative sigma factor SigG: its operon and regulon. Tuberculosis (Edinburgh, Scotland) 93: 482-491.

Gebhard S, Humpel A, McLellan AD & Cook GM (2008) The alternative sigma factor SigF of Mycobacterium smegmatis is required for survival of heat shock, acidic pH and oxidative stress. Microbiology (Reading, England) 154: 2786-2795.

Giangrossi M, Prosseda G, Tran CN, Brandi A, Colonna B & Falconi M (2010) A novel antisense RNA regulates at transcriptional level the virulence gene icsA of Shigella flexneri. Nucleic Acids Res 38: 3362-3375.

Gill EE, Chan LS, Winsor GL, et al. (2018) High-throughput detection of RNA processing in bacteria. BMC Genomics 19: 223.

Gomes AL, Abeel T, Peterson M, Azizi E, Lyubetskaya A, Carvalho L & Galagan J (2014) Decoding ChIP-seq with a double-binding signal refines binding peaks to single-nucleotides and predicts cooperative interaction. Genome research 24: 1686-1697.

Gray TA, Palumbo MJ & Derbyshire KM (2013) Draft Genome Sequence of MKD8, a Conjugal Recipient Mycobacterium smegmatis Strain. Genome announcements 1: e0014813.

Griffin JE, Gawronski JD, Dejesus MA, Ioerger TR, Akerley BJ & Sassetti CM (2011) High-resolution phenotypic profiling defines genes essential for mycobacterial growth and cholesterol catabolism. PLoS pathogens 7: e1002251.

Guell M, van Noort V, Yus E, et al. (2009) Transcriptome complexity in a genome-reduced bacterium. Science (New York, NY) 326: 1268-1271.

Gutgsell NS & Jain C (2010) Coordinated regulation of 23S rRNA maturation in Escherichia coli. J Bacteriol 192: 1405-1409.

Hartkoorn RC, Sala C, Magnet SJ, Chen JM, Pojer F & Cole ST (2010) Sigma factor F does not prevent rifampin inhibition of RNA polymerase or cause rifampin tolerance in Mycobacterium tuberculosis. J Bacteriol 192: 5472-5479.

Hayashi JM, Richardson K, Melzer ES, Sandler SJ, Aldridge BB, Siegrist MS & Morita YS (2018) Stress-induced reorganization of the mycobacterial membrane domain. mBio 9: e01823-01817.

Heidrich N, Bauriedl S, Barquist L, Li L, Schoen C & Vogel J (2017) The primary transcriptome of Neisseria meningitidis and its interaction with the RNA chaperone Hfq. Nucleic acids research 45: 6147-6167.

Heroven A, Sest M, Pisano F, Scheb-Wetzel M, Böhme K, Klein J, Münch R, Schomburg D & Dersch P (2012) Crp induces switching of the CsrB and CsrC RNAs in Yersinia pseudotuberculosis and links nutritional status to virulence. Frontiers in cellular and infection microbiology 2: 158.

Holmqvist E, Wright PR, Li L, Bischler T, Barquist L, Reinhardt R, Backofen R & Vogel J (2016) Global RNA recognition patterns of post-transcriptional regulators Hfq and CsrA revealed by UV crosslinking in vivo. 35: 991-1011.

Honaker RW, Leistikow RL, Bartek IL & Voskuil MI (2009) Unique roles of DosT and DosS in DosR regulon induction and Mycobacterium tuberculosis dormancy. Infection and immunity 77: 3258-3263.

Humpel A, Gebhard S, Cook GM & Berney M (2010) The SigF regulon in Mycobacterium smegmatis reveals roles in adaptation to stationary phase, heat, and oxidative stress. J Bacteriol 192: 2491-2502.

Ignatov DV, Salina EG, Fursov MV, Skvortsov TA, Azhikina TL & Kaprelyants AS (2015) Dormant non-culturable Mycobacterium tuberculosis retains stable low-abundant mRNA. BMC Genomics 16: 954.

Innocenti N, Golumbeanu M, Fouquier d’Herouel A, et al. (2015) Whole-genome mapping of 5′ RNA ends in bacteria by tagged sequencing: a comprehensive view in Enterococcus faecalis. RNA (New York, NY) 21: 1018-1030.

Iona E, Pardini M, Mustazzolu A, Piccaro G, Nisini R, Fattorini L & Giannoni F (2016) Mycobacterium tuberculosis gene expression at different stages of hypoxia-induced dormancy and upon resuscitation. Journal of microbiology (Seoul, Korea) 54: 565-572.

Jarmer H, Larsen TS, Krogh A, Saxild HH, Brunak S & Knudsen S (2001) Sigma A recognition sites in the Bacillus subtilis genome. Microbiology (Reading, England) 147: 2417-2424.

Jurėnaitė M, Markuckas A & Sužiedėlienė E (2013) Identification and characterization of type II toxin-antitoxin systems in the opportunistic pathogen Acinetobacter baumannii. Journal of bacteriology JB. 00237-00213.

Kawano M, Aravind L & Storz G (2007) An antisense RNA controls synthesis of an SOS-induced toxin evolved from an antitoxin. Mol Microbiol 64: 738-754.

Kovacs L, Csanadi A, Megyeri K, Kaberdin VR & Miczak A (2005) Mycobacterial RNase E-associated proteins. Microbiology and immunology 49: 1003-1007.

Kulesekara H, Lee V, Brencic A, Liberati N, Urbach J, Miyata S, Lee DG, Neely AN, Hyodo M & Hayakawa Y (2006) Analysis of Pseudomonas aeruginosa diguanylate cyclases and phosphodiesterases reveals a role for bis-(3′-5′)-cyclic-GMP in virulence. Proceedings of the National Academy of Sciences 103: 2839-2844.

Lee JH, Karakousis PC & Bishai WR (2008) Roles of SigB and SigF in the Mycobacterium tuberculosis sigma factor network. J Bacteriol 190: 699-707.

Lee JH, Geiman DE & Bishai WR (2008) Role of stress response sigma factor SigG in Mycobacterium tuberculosis. J Bacteriol 190: 1128-1133.

Leistikow RL, Morton RA, Bartek IL, Frimpong I, Wagner K & Voskuil MI (2010) The Mycobacterium tuberculosis DosR regulon assists in metabolic homeostasis and enables rapid recovery from nonrespiring dormancy. J Bacteriol 192: 1662-1670.

Lewis DE & Adhya S (2004) Axiom of determining transcription start points by RNA polymerase in Escherichia coli. Mol Microbiol 54: 692-701.

Li H & Durbin R (2009) Fast and accurate short read alignment with Burrows-Wheeler transform. Bioinformatics (Oxford, England) 25: 1754-1760.

Li X, Mei H, Chen F, Tang Q, Yu Z, Cao X, Andongma BT, Chou S-H & He J (2017) Transcriptome landscape of Mycobacterium smegmatis. Frontiers in microbiology 8: 2505.

Li Z & Deutscher MP (1996) Maturation pathways for E. coli tRNA precursors: a random multienzyme process in vivo. Cell 86: 503-512.

Llorens-Rico V & Cano J (2016) Bacterial antisense RNAs are mainly the product of transcriptional noise. 2: e1501363.

Love MI, Huber W & Anders S (2014) Moderated estimation of fold change and dispersion for RNA-seq data with DESeq2. Genome Biology 15: 550.

Lun DS, Sherrid A, Weiner B, Sherman DR & Galagan JE (2009) A blind deconvolution approach to highresolution mapping of transcription factor binding sites from ChIP-seq data. Genome Biol 10: R142.

Mackie GA (2013) RNase E: at the interface of bacterial RNA processing and decay. Nature reviews Microbiology 11: 45-57.

McKenzie JL, Robson J, Berney M, Smith TC, Ruthe A, Gardner PP, Arcus VL & Cook GM (2012) A VapBC toxin-antitoxin module is a posttranscriptional regulator of metabolic flux in mycobacteria. J Bacteriol 194: 2189-2204.

Mendoza-Vargas A, Olvera L, Olvera M, et al. (2009) Genome-wide identification of transcription start sites, promoters and transcription factor binding sites in E. coli. PLoS One 4: e7526.

Michele TM, Ko C & Bishai WR (1999) Exposure to antibiotics induces expression of the Mycobacterium tuberculosis sigF gene: implications for chemotherapy against mycobacterial persistors. Antimicrobial agents and chemotherapy 43: 218-225.

Milano A, Forti F, Sala C, Riccardi G & Ghisotti D (2001) Transcriptional regulation of furA and katG upon oxidative stress in Mycobacterium smegmatis. J Bacteriol 183: 6801-6806.

Mitschke J, Georg J, Scholz I, Sharma CM, Dienst D, Bantscheff J, Voß B, Steglich C, Wilde A & Vogel J (2011) An experimentally anchored map of transcriptional start sites in the model cyanobacterium Synechocystis sp. PCC6803. Proceedings of the National Academy of Sciences 108: 2124-2129.

Moores A, Riesco AB, Schwenk S & Arnvig KB (2017) Expression, maturation and turnover of DrrS, an unusually stable, DosR regulated small RNA in Mycobacterium tuberculosis. 12: e0174079.

Morita T, Maki K & Aiba H (2005) RNase E-based ribonucleoprotein complexes: mechanical basis of mRNA destabilization mediated by bacterial noncoding RNAs. Genes & development 19: 2176-2186.

Mraheil MA, Billion A, Mohamed W, Mukherjee K, Kuenne C, Pischimarov J, Krawitz C, Retey J, Hartsch T & Chakraborty T (2011) The intracellular sRNA transcriptome of Listeria monocytogenes during growth in macrophages. Nucleic acids research 39: 4235-4248.

Newton-Foot M & Gey van Pittius NC (2013) The complex architecture of mycobacterial promoters. Tuberculosis (Edinburgh, Scotland) 93: 60-74.

Novichkov PS, Kazakov AE, Ravcheev DA, et al. (2013) RegPrecise 3.0‐‐a resource for genome-scale exploration of transcriptional regulation in bacteria. BMC Genomics 14: 745.

O’Toole R, Smeulders MJ, Blokpoel MC, Kay EJ, Lougheed K & Williams HD (2003) A two-component regulator of universal stress protein expression and adaptation to oxygen starvation in Mycobacterium smegmatis. Journal of bacteriology 185: 1543-1554.

Paletta JL & Ohman DE (2012) Evidence for two promoters internal to the alginate biosynthesis operon in Pseudomonas aeruginosa. Current microbiology 65: 770-775.

Park HD, Guinn KM, Harrell MI, Liao R, Voskuil MI, Tompa M, Schoolnik GK & Sherman DR (2003) Rv3133c/dosR is a transcription factor that mediates the hypoxic response of Mycobacterium tuberculosis. Molecular microbiology 48: 833-843.

Pecsi I, Hards K, Ekanayaka N, Berney M, Hartman T, Jacobs WR & Cook GM (2014) Essentiality of succinate dehydrogenase in Mycobacterium smegmatis and its role in the generation of the membrane potential under hypoxia. MBio 5: e01093-01014.

Potgieter MG, Nakedi KC, Ambler JM, Nel AJ, Garnett S, Soares NC, Mulder N & Blackburn JM (2016) Proteogenomic Analysis of Mycobacterium smegmatis using high resolution mass spectrometry. Frontiers in microbiology 7: 427.

Prasanna AN & Mehra S (2013) Comparative phylogenomics of pathogenic and non-pathogenic mycobacterium. PLoS One 8: e71248.

Raju RM, Unnikrishnan M, Rubin DH, Krishnamoorthy V, Kandror O, Akopian TN, Goldberg AL & Rubin EJ (2012) Mycobacterium tuberculosis ClpP1 and ClpP2 function together in protein degradation and are required for viability in vitro and during infection. PLoS pathogens 8: e1002511.

Ramachandran VK, Shearer N & Thompson A (2014) The primary transcriptome of Salmonella enterica serovar Typhimurium and its dependence on ppGpp during late stationary phase. PLoS One 9: e92690.

Raman S, Hazra R, Dascher CC & Husson RN (2004) Transcription regulation by the Mycobacterium tuberculosis alternative sigma factor SigD and its role in virulence. J Bacteriol 186: 6605-6616.

Raman S, Song T, Puyang X, Bardarov S, Jacobs WR, Jr. & Husson RN (2001) The alternative sigma factor SigH regulates major components of oxidative and heat stress responses in Mycobacterium tuberculosis. J Bacteriol 183: 6119-6125.

Rigel NW, Gibbons HS, McCann JR, McDonough JA, Kurtz S & Braunstein M (2009) The accessory SecA2 system of mycobacteria requires ATP binding and the canonical SecA1. Journal of Biological Chemistry 284: 9927-9936.

Roberts DM, Liao RP, Wisedchaisri G, Hol WG & Sherman DR (2004) Two sensor kinases contribute to the hypoxic response of Mycobacterium tuberculosis. The Journal of biological chemistry 279: 23082-23087.

Robson J, McKenzie JL, Cursons R, Cook GM & Arcus VL (2009) The vapBC operon from Mycobacterium smegmatis is an autoregulated toxin-antitoxin module that controls growth via inhibition of translation. Journal of molecular biology 390: 353-367.

Rodrigue S, Brodeur J, Jacques PE, Gervais AL, Brzezinski R & Gaudreau L (2007) Identification of mycobacterial sigma factor binding sites by chromatin immunoprecipitation assays. J Bacteriol 189: 1505-1513.

Rosenkrands I, Slayden RA, Crawford J, Aagaard C, Clifton III E & Andersen P (2002) Hypoxic response of Mycobacterium tuberculosis studied by metabolic labeling and proteome analysis of cellular and extracellular proteins. Journal of bacteriology 184: 3485-3491.

Rustad TR, Harrell MI, Liao R & Sherman DR (2008) The enduring hypoxic response of Mycobacterium tuberculosis. PloS one 3: e1502.

Rustad TR, Minch KJ, Brabant W, Winkler JK, Reiss DJ, Baliga NS & Sherman DR (2013) Global analysis of mRNA stability in Mycobacterium tuberculosis. Nucleic Acids Res 41: 509-517.

Sala C, Forti F, Magnoni F & Ghisotti D (2008) The katG mRNA of Mycobacterium tuberculosis and Mycobacterium smegmatis is processed at its 5′ end and is stabilized by both a polypurine sequence and translation initiation. BMC molecular biology 9: 33.

Sass AM, Van Acker H, Förstner KU, Van Nieuwerburgh F, Deforce D, Vogel J & Coenye T (2015) Genomewide transcription start site profiling in biofilm-grown Burkholderia cenocepacia J2315. BMC genomics 16:775.

Sassetti CM & Rubin EJ (2003) Genetic requirements for mycobacterial survival during infection. Proceedings of the National Academy of Sciences of the United States of America 100: 12989-12994.

Sassetti CM, Boyd DH & Rubin EJ (2003) Genes required for mycobacterial growth defined by high density mutagenesis. Mol Microbiol 48: 77-84.

Schifano JM, Edifor R, Sharp JD, Ouyang M, Konkimalla A, Husson RN & Woychik NA (2013) Mycobacterial toxin MazF-mt6 inhibits translation through cleavage of 23S rRNA at the ribosomal A site. Proceedings of the National Academy of Sciences 110: 8501-8506.

Schlüter J-P, Reinkensmeier J, Barnett MJ, Lang C, Krol E, Giegerich R, Long SR & Becker A (2013) Global mapping of transcription start sites and promoter motifs in the symbiotic α-proteobacterium Sinorhizobium meliloti 1021. BMC genomics 14: 156.

Shao W, Price MN, Deutschbauer AM, Romine MF & Arkin AP (2014) Conservation of transcription start sites within genes across a bacterial genus. MBio 5: e01398-01314.

Sharma CM, Hoffmann S, Darfeuille F, et al. (2010) The primary transcriptome of the major human pathogen Helicobacter pylori. Nature 464: 250-255.

Shell SS, Chase MR, Ioerger TR & Fortune SM (2015) RNA sequencing for transcript 5′-end mapping in mycobacteria. Methods in molecular biology (Clifton, NJ) 1285: 31-45.

Shell SS, Wang J, Lapierre P, Mir M, Chase MR, Pyle MM, Gawande R, Ahmad R, Sarracino DA & Ioerger TR (2015) Leaderless transcripts and small proteins are common features of the mycobacterial translational landscape. PLoS genetics 11: e1005641.

Singh AK & Singh BN (2008) Conservation of sigma F in mycobacteria and its expression in Mycobacterium smegmatis. Current microbiology 56: 574-580.

Singh AK, Dutta D, Singh V, Srivastava V, Biswas RK & Singh BN (2015) Characterization of Mycobacterium smegmatis sigF mutant and its regulon: overexpression of SigF antagonist (MSMEG_1803) in M. smegmatis mimics sigF mutant phenotype, loss of pigmentation, and sensitivity to oxidative stress. MicrobiologyOpen 4: 896-916.

Skliarova SA, Kreneva RA, Perumov DA & Mironov AS (2012) [The characterization of internal promoters in the Bacillus subtilis riboflavin biosynthesis operon]. Genetika 48: 1133-1141.

Song T, Song SE, Raman S, Anaya M & Husson RN (2008) Critical role of a single position in the -35 element for promoter recognition by Mycobacterium tuberculosis SigE and SigH. J Bacteriol 190: 2227-2230.

Sun R, Converse PJ, Ko C, Tyagi S, Morrison NE & Bishai WR (2004) Mycobacterium tuberculosis ECF sigma factor sigC is required for lethality in mice and for the conditional expression of a defined gene set. Mol Microbiol 52: 25-38.

Taverniti V, Forti F, Ghisotti D & Putzer H (2011) Mycobacterium smegmatis RNase J is a 5′-3′ exo-/endoribonuclease and both RNase J and RNase E are involved in ribosomal RNA maturation. Mol Microbiol 82: 1260-1276.

Thomason MK, Bischler T, Eisenbart SK, Förstner KU, Zhang A, Herbig A, Nieselt K, Sharma CM & Storz G (2015) Global transcriptional start site mapping using differential RNA sequencing reveals novel antisense RNAs in Escherichia coli. Journal of bacteriology 197: 18-28.

Trauner A, Lougheed KE, Bennett MH, Hingley-Wilson SM & Williams HD (2012) The dormancy regulator DosR controls ribosome stability in hypoxic mycobacteria. Journal of Biological Chemistry jbc. M112. 364851.

Veyrier F, Said-Salim B & Behr MA (2008) Evolution of the mycobacterial SigK regulon. J Bacteriol 190: 1891-1899.

Warner DF, Savvi S, Mizrahi V & Dawes SS (2007) A riboswitch regulates expression of the coenzyme B12-independent methionine synthase in Mycobacterium tuberculosis: implications for differential methionine synthase function in strains H37Rv and CDC1551. J Bacteriol 189: 3655-3659.

Wayne LG & Hayes LG (1996) An in vitro model for sequential study of shiftdown of Mycobacterium tuberculosis through two stages of nonreplicating persistence. Infection and immunity 64: 2062-2069.

Winther K, Tree JJ, Tollervey D & Gerdes K (2016) VapCs of Mycobacterium tuberculosis cleave RNAs essential for translation. Nucleic Acids Res 44: 9860-9871.

Wittchen M, Busche T, Gaspar AH, Lee JH, Ton-That H, Kalinowski J & Tauch A (2018) Transcriptome sequencing of the human pathogen Corynebacterium diphtheriae NCTC 13129 provides detailed insights into its transcriptional landscape and into DtxR-mediated transcriptional regulation. BMC Genomics 19: 82.

Wu M-L, Gengenbacher M & Dick T (2016) Mild nutrient starvation triggers the development of a smallcell survival morphotype in mycobacteria. Frontiers in microbiology 7: 947.

Yang H, Sha W, Liu Z, et al. (2018) Lysine acetylation of DosR regulates the hypoxia response of Mycobacterium tuberculosis. Emerging microbes & infections 7: 34.

Zeller ME, Csanadi A, Miczak A, Rose T, Bizebard T & Kaberdin VR (2007) Quaternary structure and biochemical properties of mycobacterial RNase E/G. The Biochemical journal 403: 207-215.

Zhu Y, Mao C, Ge X, Wang Z, Lu P, Zhang Y, Chen S & Hu Y (2017) Characterization of a Minimal Type of Promoter Containing the -10 Element and a Guanine at the -14 or -13 Position in Mycobacteria. J Bacteriol 199.

Zhukova A, Fernandes LG, Hugon P, et al. (2017) Genome-Wide Transcriptional Start Site Mapping and sRNA Identification in the Pathogen Leptospira interrogans. Front Cell Infect Microbiol 7: 10.

